# Genetic expression of 4E-BP1 in juvenile mice alleviates mTOR-induced neuronal dysfunction and epilepsy

**DOI:** 10.1101/2021.06.04.445850

**Authors:** Lena H. Nguyen, Youfen Xu, Travorn Mahadeo, Longbo Zhang, Tiffany V. Lin, Heather A. Born, Anne E. Anderson, Angélique Bordey

**Affiliations:** Departments of Neurosurgery, Yale University School of Medicine, New Haven, CT 06510, USA; Departments of Cellular & Molecular Physiology, Yale University School of Medicine, New Haven, CT 06510, USA; Departments of Cain Foundation Laboratories, Jan and Dan Duncan Neurological Research Institute at Texas Children’s Hospital, Houston, TX 77030, USA; Department of Pediatrics, Baylor College of Medicine, Houston, TX 77030, USA

**Keywords:** mTOR, 4E-BP1, malformation of cortical development, epilepsy, neurodevelopmental disorders, neuronal excitability, HCN channels, EEG, spectral analysis, *in utero* electroporation

## Abstract

Hyperactivation of mTOR signaling during fetal neurodevelopment alters neuron structure and function, leading to focal malformation of cortical development (FMCD) and intractable epilepsy. Recent evidence suggests increased cap-dependent translation downstream of mTOR contributes to FMCD formation and seizures. However, whether reducing overactive translation once the developmental pathologies are established reverses neuronal abnormalities and seizures is unknown. Here, we found that the translational repressor 4E-BP1, which is inactivated by mTOR-mediated phosphorylation, is hyperphosphorylated in patient FMCD tissue and in a mouse model of FMCD. Expressing constitutive active 4E-BP1 to repress aberrant translation in juvenile mice with FMCD reduced neuronal cytomegaly and corrected several electrophysiological alterations, including depolarized resting membrane potential, irregular firing pattern, and aberrant HCN4 channel expression. This was accompanied by improved cortical spectral activity and decreased seizures. Although mTOR controls multiple pathways, our study shows that targeting 4E-BP1-mediated translation alone is sufficient to alleviate neuronal dysfunction and ongoing epilepsy.

## INTRODUCTION

Hyperactivation of mechanistic target of rapamycin (mTOR) signaling causes a spectrum of neurodevelopmental disorders associated with focal malformation of cortical development (FMCD), including tuberous sclerosis complex (TSC), focal cortical dysplasia type II (FCDII), and hemimegalencephaly (HME) (Crino, 2015). These disorders result from somatic or germline mutations in mTOR complex 1 (mTORC1) pathway genes and are the most common causes for intractable epilepsy in children (Kwiatkowski, 2003; Marsan and Baulac, 2018; Muhlebner et al., 2019). FMCDs in TSC, FCDII, and HME share core histopathological features characterized by cortical mislamination, neuronal heterotopia, and the presence of dysmorphic cytomegalic neurons with enhanced mTORC1 activation (Lim and Crino, 2013). Numerous studies in mouse models of FMCD have shown that treatment with the mTORC1 inhibitor, rapamycin, prevents FMCD formation and reduces seizures (Ostendorf and Wong, 2015; Nguyen and Bordey, 2021). These findings emphasize the central role of mTORC1 dysregulation in the etiology of these disorders and identifying the downstream mechanisms of mTORC1 that lead to FMCD and epilepsy is an active area of investigation.

The mTORC1 pathway is a key regulator of cell growth that controls multiple cellular processes, including cap-dependent translation, autophagy, lysosome biogenesis, and lipid synthesis, among others (Lipton and Sahin, 2014; Saxton and Sabatini, 2017). Hyperactive mTORC1 signaling is thought to alter neuron structure and function, leading to pro-epileptogenic circuit formation and epilepsy (Lasarge and Danzer, 2014). Despite that numerous cellular processes are dysregulated under hyperactive mTORC1 conditions, it has been reported that correcting one of these functions, in particular cap-dependent translation, during fetal neurodevelopment is sufficient to reduce seizures in a mouse model of FMCD (Kim et al., 2019). Consistent with this finding, normalizing translation before birth also rescued characteristic FMCD cytoarchitectural pathology, including neuronal misplacement, soma hypertrophy, and dendrite and axon overgrowth (Gong et al., 2015; Lin et al., 2016; Kim et al., 2019).

Although these data suggest that neuronal misplacement and dysmorphogenesis contribute to seizures, other neuronal alterations that promote epilepsy have been reported in FMCD. In particular, we previously found that dysmorphic cytomegalic neurons in TSC and FCDII patients and FMCD pyramidal neurons (modeling human dysmorphic neurons) in mice display aberrant expression of hyperpolarization-activated cyclic nucleotide-gated-isoform 4 (HCN4) channels (Hsieh et al., 2020). The abnormal presence of these channels confers a unique mode of excitability in FMCD neurons that is dependent on intracellular cyclic AMP (cAMP) levels. Silencing HCN4 channel activity before birth prevented seizures in mice with FMCD, supporting their role in seizure generation. Further, the aberrant HCN4 channel expression was prevented by rapamycin treatment, suggesting that this is an mTORC1-dependent process. However, it is unknown whether normalizing cap-dependent translation can rescue aberrant HCN4 channel expression and restore neuron excitability. Importantly, it is unknown whether modifying dysregulated translation beyond the prenatal period, once FMCD is formed and epilepsy is established, can reverse neuronal alterations and alleviate seizures. Addressing these issues is crucial with regards to therapeutics since these neurodevelopmental disorders are predominantly diagnosed during childhood, when the patients present with symptoms (Staley et al., 2011; Kabat and Krol, 2012; Ebrahimi-Fakhari et al., 2018; Racz et al., 2018).

Here, we investigated whether targeting aberrant translation once the mTORC1-induced developmental pathologies are established reverses neuronal dysfunction and seizures in a mouse model of FMCD. mTORC1 promotes cap-dependent translation by directly phosphorylating and inactivating the translational repressor, eukaryotic translation initiation factor 4E (eIF4E)-binding protein (4E-BP). Phosphorylated 4E-BPs dissociate from eIF4E, allowing eIF4E to interact with other eukaryotic initiation factors to form the eIF4F complex and initiate translation (Richter and Sonenberg, 2005; Ma and Blenis, 2009). Thus, mTORC1 hyperactivation leads 4E-BP hyperphosphorylation and overactive translation due to decreased translational repression (Lin et al., 2016). Consistent with this, we found increased levels of phosphorylated 4E-BP1 in brain tissue from TSC and FCDII patients and mice with FMCD. To compensate for hyperphosphorylated 4E-BP1 in mice, we expressed a constitutive active form of 4E-BP1 (4E-BP1^CA^) which resists phosphorylation and inactivation by mTORC1. By employing *in utero* electroporation to target specific neuronal populations and tamoxifen-inducible conditional CA vectors to attain temporal control of gene expression, we achieved timed expression of 4E-BP1^CA^ in FMCD neurons in juvenile mice. Subsequent neurophysiological analysis by patch clamp recording and video-electroencephalography (vEEG) showed that 4E-BP1^CA^ expression corrected several electrophysiological alterations, including aberrant HCN4 channel expression, normalized cortical spectral activity, and decreased seizures. Collectively, these findings support targeting 4E-BP-mediated translation as a strategy to treat epilepsy and underlying neuronal aberrancies in FMCD.

## RESULTS

### 4E-BP1 is hyperphosphorylated in human FMCD tissue and in the Rheb^CA^ mouse model of FMCD

Activated mTORC1 phosphorylates and inhibits 4E-BP repressor activity to promote cap-dependent translation, and conversely, inactivated mTORC1 leads to dephosphorylated 4E-BPs and repressed translation (Richter and Sonenberg, 2005; Ma and Blenis, 2009) (**Figure 1A**). To determine the phosphorylation states of 4E-BPs in human FMCD, we performed immunofluorescence staining for phospho (p)-4E-BP1 in resected FMCD tissue samples from individuals with TSC or FCDII who underwent epilepsy surgery. We co-stained with the neurofilament marker SMI-311 to identify the dysmorphic cytomegalic neurons. These neurons have an abnormal cytoplasmic accumulation of neurofilament proteins that result in intense neurofilament labeling (Sisodiya et al., 2009; Blumcke et al., 2011; Sousa et al., 2018). We found marked p-4E-BP1 immunoreactivity in SMI-311+ dysmorphic neurons, whereas low or no p-4E-BP1 immunoreactivity was detected in surrounding SMI-311-cells in all samples (**Figure 1B**). Increased p-4E-BP1 immunoreactivity was also observed in additional FCDII samples by immunohistochemical staining (**Supplemental Figure 1A**). These results demonstrate the occurrence of 4E-BP1 hyperphosphorylation in human FMCD dysmorphic neurons and suggest the presence of overactive translation in these cells.

**Figure 1:**
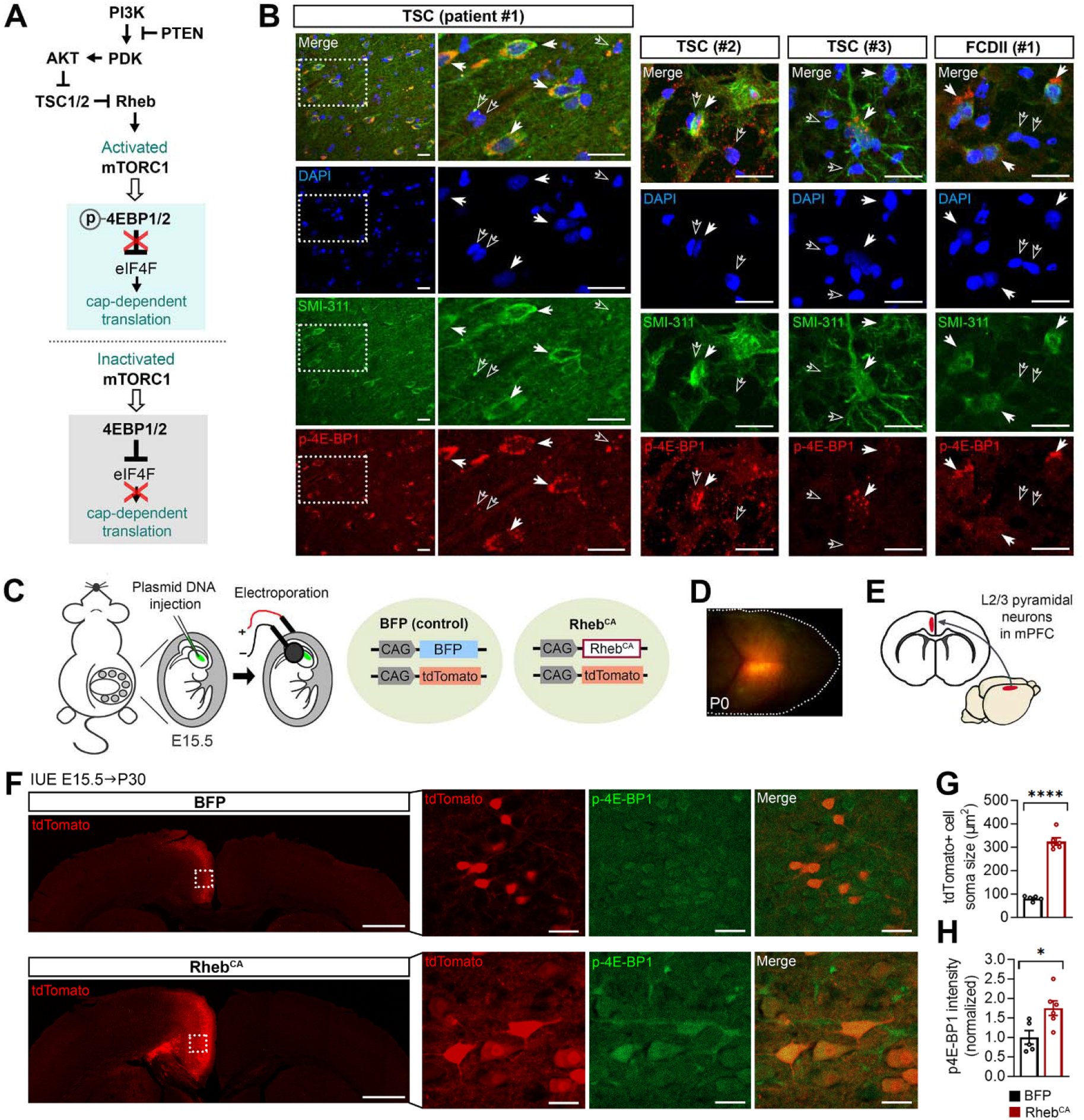
4E-BP1 is hyperphosphorylated in human FMCD tissue and in the Rheb^CA^ mouse model of FMCD. **(A)** Diagram of the PI3K-mTORC1 pathway and downstream regulation of cap-dependent translation via 4E-BP1/2. Activated mTORC1 phosphorylates and inhibits 4E-BP1/2, which disinhibits the eIF4F complex (via release of eIF4E) and promotes cap-dependent translation. Conversely, inactivated mTORC1 disinhibits 4E-BP1/2, which inhibits 4E-BP1/2 and blocks cap-dependent translation. **(B)** Representative images of DAPI (blue), SMI-311 (green), and phospho-4E-BP1 (p-4E-BP1, red) staining in resected brain tissue samples from 3 individuals with TSC and 1 individual with FCDII who underwent epilepsy surgery. For individual TSC #1, low magnification images are shown on the left and high magnification images are shown on the right. White squares denote magnified areas. For all images, filled arrows point to SMI-311+ dysmorphic neurons with high p-4E-BP1 immunoreactivity. Unfilled arrows point to surrounding SMI-311-cells with low or no p-4E-BP1 immunoreactivity. Note that DAPI-stained nuclei in SMI-311+ cells appear fainter and are often enlarged compared to SMI-311-cells. Scale bars=25 μm. **(C)** Diagram of IUE and plasmids used to generate Rheb^CA^ mice with FMCD. Mouse embryos were electroporated with a BFP (control) or Rheb^CA^ plasmid at E15.5, targeting radial glia generating pyramidal neurons destined to L2/3 in the mPFC. tdTomato was co-electroporated in both conditions to label the targeted neurons. **(D)** Image of a P0 pup head showing tdTomato fluorescence in the targeted region following electroporation at E15.5. **(E)** Diagram showing the targeted region (red) in an adult mouse brain and the corresponding area in coronal view. **(F)** Representative images of tdTomato+ cells (red) and p-4E-BP1 staining (green, pseudocolored) in coronal cortical sections from P30 BFP control and Rheb^CA^ mice. Low magnification tile scan images of tdTomato are shown on the left. tdTomato+ cells are strictly found in L2/3 in BFP control mice and misplaced across the cortical layers in Rheb^CA^ mice. High magnification images of tdTomato+ cells and p-4E-BP1 staining are shown on the right. tdTomato+ cells display basal level p-4E-BP1 immunoreactivity in BFP control mice and high p-4E-BP1 immunoreactivity in Rheb^CA^ mice. White squares denote the magnified areas. Scale bars=1000 μm (left), 25 μm (right). **(G)** Quantification of tdTomato+ cell soma size. n=5 BFP, 6 Rheb^CA^ mice; each data point represents averaged values from 22-30 cells per animal. Data were analyzed by unpaired t-test; ****p<0.0001. **(H)** Quantification of p-4E-BP1 intensity in tdTomato+ cells. n=5 BFP, 6 Rheb^CA^ mice; each data point represents averaged values from 22-30 cells per animal. Data were normalized to the mean control and analyzed by unpaired t-test; *p=0.0186. Error bars are ± SEM.

To model FMCD in mice, we expressed a constitutive active form of the canonical mTORC1 activator, Rheb (Rheb^CA^), in developing mouse embryos using IUE, as previously described (Hsieh et al., 2016; Lin et al., 2016; Nguyen et al., 2019; Hsieh et al., 2020; Zhang et al., 2020b). Mice were electroporated with plasmids encoding for Rheb^CA^ or BFP (as control) at embryonic day (E) 15.5, targeting radial glia that generate pyramidal neurons destined to layer (L) 2/3 in the medial prefrontal cortex (mPFC). A tdTomato reporter plasmid was co-electroporated in both conditions to label the targeted neurons (**Figure 1C-E**). Histological examination was performed in juvenile mice at postnatal day (P) 30. Consistent with previous findings, Rheb^CA^ expression resulted in neuron misplacement across the cortical layers (**Figure 1F**) and increased neuron soma size (**Figures 1F, G, Table 1**) (Hsieh et al., 2016; Lin et al., 2016; Nguyen et al., 2019; Zhang et al., 2020b). Immunostaining for p-4E-BP1 revealed significantly increased p-4E-BP1 levels in Rheb^CA^ neurons compared to control neurons (**Figure 1F, H, Table 1**). Similar results were also found when stained with a p-4E-BP1/2/3 antibody that recognizes all 4E-BP isoforms at P30 and P165 (**Supplemental Figure 1B-E, Table S1**). Thus, dysregulation of 4E-BP1 occurs in both human and mouse FMCD.

**Table 1:** Table of main results and statistics.

| Figure | Statistical test, results | Group mean, sample size |
| --- | --- | --- |
| **1G**<br>Cell soma size<br>(P30) | **Unpaired t-test (two-tailed)**<br>p value <0.0001<br>t, df t=13.90, df=9 | **BFP**<br>Mean ± SD 81.3 ± 10.5<br>Sample size (n) 5 mice<br>**Rheb<sup>CA</sup>**<br>Mean ± SD 324.7 ± 37.7<br>Sample size (n) 6 mice |
| **1H**<br>p-4E-BP1 intensity<br>(P30) | **Unpaired t-test (two-tailed)**<br>p value 0.0186<br>t, df t=2.866, df=9 | **BFP**<br>Mean ± SD 1.00 ± 0.39<br>Sample size (n) 5 mice<br>**Rheb<sup>CA</sup>**<br>Mean ± SD 1.75 ± 0.46<br>Sample size (n) 6 mice |
| **2C**<br>Membrane<br>capacitance | **One-way ANOVA**<br>p value <0.0001<br>F (DFn, DFd) F (2, 45) = 25.36 | **BFP+cGFP**<br>Mean ± SD 150.7 ± 39.8<br>Sample size (n) 14 neurons<br>**Rheb<sup>CA</sup>+cGFP**<br>Mean ± SD 402.9 ± 131.4<br>Sample size (n) 16 neurons<br>**Rheb<sup>CA</sup>+c4E-BP1<sup>CA</sup>**<br>Mean ± SD 317.0 ± 94.6<br>Sample size (n) 18 neurons |
| **2D**<br>Conductance | **One-way ANOVA**<br>p value <0.0001<br>F (DFn, DFd) F (2, 44) = 11.82 | **BFP+cGFP**<br>Mean ± SD 8.7 ± 2.3<br>Sample size (n) 14 neurons<br>**Rheb<sup>CA</sup>+cGFP**<br>Mean ± SD 16.6 ± 5.4<br>Sample size (n) 15 neurons<br>**Rheb<sup>CA</sup>+c4E-BP1<sup>CA</sup>**<br>Mean ± SD 14.9 ± 5.1<br>Sample size (n) 18 neurons |
| **2E**<br>RMP | **One-way ANOVA**<br>p value 0.0031<br>F (DFn, DFd) F (2, 45) = 6.596 | **BFP+cGFP**<br>Mean ± SD -73.1 ± 5.6<br>Sample size (n) 14 neurons<br>**Rheb<sup>CA</sup>+cGFP**<br>Mean ± SD -65.0 ± 7.8<br>Sample size (n) 16 neurons<br>**Rheb<sup>CA</sup>+c4E-BP1<sup>CA</sup>**<br>Mean ± SD -70.4 ± 5.3<br>Sample size (n) 18 neurons |
| **2G**<br>IV curve (of I<sub>ss</sub>) | **Mixed-effects ANOVA**<br>**Voltage step**<br>p value <0.0001<br>F (DFn, DFd) F (1.337, 60.02) = 179.0<br>**Group**<br>p value 0.0002<br>F (DFn, DFd) F (2, 45) = 10.23<br>**Voltage step x group**<br>p value <0.0001<br>F (DFn, DFd) F (18, 404) = 9.798 | **BFP+cGFP**<br>Sample size (n) 14 neurons<br>**Rheb<sup>CA</sup>+cGFP**<br>Sample size (n) 16 neurons<br>**Rheb<sup>CA</sup>+c4E-BP1<sup>CA</sup>**<br>Sample size (n) 18 neurons |
| **2H**<br>IV curve (of $I_h$) | **Mixed-effects ANOVA**<br>**Voltage step**<br>p value <0.0001<br>F (DFn, DFd) F (1.131, 47.90) = 33.60<br>**Group**<br>p value 0.0004<br>F (DFn, DFd) F (2, 45) = 9.256<br>**Voltage step x group**<br>p value <0.0001<br>F (DFn, DFd) F (18, 381) = 7.404 | **BFP+cGFP**<br>Sample size (n) 14 neurons<br>***Rheb*<sup>CA</sup>+cGFP**<br>Sample size (n) 16 neurons<br>***Rheb*<sup>CA</sup>+c4E-BP1<sup>CA</sup>**<br>Sample size (n) 18 neurons |
| **2I**<br>$I_h$ at -90 mV | **One-way ANOVA**<br>p value 0.0002<br>F (DFn, DFd) F (2, 44) = 10.59 | **BFP+cGFP**<br>Mean ± SD -12.00 ± 6.9<br>Sample size (n) 14 neurons<br>***Rheb*<sup>CA</sup>+cGFP**<br>Mean ± SD -102.6 ± 96.9<br>Sample size (n) 16 neurons<br>***Rheb*<sup>CA</sup>+c4E-BP1<sup>CA</sup>**<br>Mean ± SD -27.42 ± 30.8<br>Sample size (n) 17 neurons |
| **2J**<br>$I_h$ amplitude vs. RMP | **Pearson product-moment correlation**<br>p value 0.0021<br>Pearson r -0.4371 | Number of XY pairs 47 |
| **2M**<br>Sag ratio | **One-way ANOVA**<br>p value 0.0016<br>F (DFn, DFd) F (2, 40) = 7.566 | **BFP+cGFP**<br>Mean ± SD 3.4 ± 3.2<br>Sample size (n) 10 neurons<br>***Rheb*<sup>CA</sup>+cGFP**<br>Mean ± SD 15.3 ± 13.8<br>Sample size (n) 15 neurons<br>***Rheb*<sup>CA</sup>+c4E-BP1<sup>CA</sup>**<br>Mean ± SD 4.9 ± 4.5<br>Sample size (n) 18 neurons |
| **2O**<br>Input-output curve | **Mixed-effects ANOVA**<br>**Current step**<br>p value <0.0001<br>F (DFn, DFd) F (1.442, 49.01) = 163.1<br>**Group**<br>p value 0.0861<br>F (DFn, DFd) F (2, 37) = 2.622<br>**Current step x group**<br>p value 0.0001<br>F (DFn, DFd) F (28, 476) = 2.376 | **BFP+cGFP**<br>Sample size (n) 10 neurons<br>***Rheb*<sup>CA</sup>+cGFP**<br>Sample size (n) 13 neurons<br>***Rheb*<sup>CA</sup>+c4E-BP1<sup>CA</sup>**<br>Sample size (n) 17 neurons |
| **3C**<br>Cell placement (P84-106) | **Unpaired t-test (two-tailed)**<br>p value 0.0533<br>t, df t=2.021, df=27 | ***Rheb*<sup>CA</sup>+cGFP**<br>Mean ± SD 34.1 ± 12.9<br>Sample size (n) 11 mice<br>***Rheb*<sup>CA</sup>+c4E-BP1<sup>CA</sup>**<br>Mean ± SD 42.4 ± 9.4<br>Sample size (n) 18 mice |
| **3D**<br>Cell soma size (P84-106) | **Unpaired t-test (two-tailed)**<br>p value <0.0001<br>t, df t=9.691, df=27 | ***Rheb*<sup>CA</sup>+cGFP**<br>Mean ± SD 464.3 ± 64.9<br>Sample size (n) 11 mice<br>***Rheb*<sup>CA</sup>+c4E-BP1<sup>CA</sup>**<br>Mean ± SD 246.1 ± 55.0<br>Sample size (n) 18 mice |
| **3F**<br>HCN4 intensity | **Two-way repeated measures ANOVA**<br>**Cortex side** | ***Rheb*<sup>CA</sup>+cGFP**<br>Mean ± SD (ipsi) 1.79 ± 0.40 |
|  | p value <0.0001<br>F (DFn, DFd) F (1, 26) = 65.51<br>**Group**<br>p value 0.001<br>F (DFn, DFd) F (1, 26) = 13.83<br>**Cortical side x group**<br>p value 0.0004<br>F (DFn, DFd) F (1, 26) = 16.55 | Mean ± SD (contra) 1.00 ± 0.17<br>Sample size (n) 11 mice<br>***Rheb<sup>CA</sup>+c4E-BP1<sup>CA</sup>***<br>Mean ± SD (ipsi) 1.19 ± 0.35<br>Mean ± SD (contra) 0.93 ± 0.14<br>Sample size (n) 17 mice |
| **4B**<br>Power spectra, light | **Two-way repeated measures ANOVA**<br>**Frequency bin**<br>p value <0.0001<br>F (DFn, DFd) F (2.32, 74.07) = 962.7<br>**Group**<br>p value 0.9157<br>F (DFn, DFd) F (2, 32) = 0.08835<br>**Frequency bin x group**<br>p value 0.0003<br>F (DFn, DFd) F (124, 1984) = 1.518 | ***GFP***<br>Sample size (n) 6 mice<br>***Rheb<sup>CA</sup>+cGFP***<br>Sample size (n) 11 mice<br>***Rheb<sup>CA</sup>+c4E-BP1<sup>CA</sup>***<br>Sample size (n) 18 mice |
| **4C**<br>Power spectra, dark | **Two-way repeated measures ANOVA**<br>**Frequency bin**<br>p value <0.0001<br>F (DFn, DFd) F (2.74, 87.51) = 504.4<br>**Group**<br>p value 0.8166<br>F (DFn, DFd) F (2, 32) = 0.2040<br>**Frequency bin x group**<br>p value 0.2840<br>F (DFn, DFd) F (124, 1984) = 1.071 | ***GFP***<br>Sample size (n) 6 mice<br>***Rheb<sup>CA</sup>+cGFP***<br>Sample size (n) 11 mice<br>***Rheb<sup>CA</sup>+c4E-BP1<sup>CA</sup>***<br>Sample size (n) 18 mice |
| **4E**<br>Bandpower, light<br>Delta | **One-way ANOVA**<br>p value 0.2874<br>F (DFn, DFd) F (2, 32) = 1.297 | ***GFP***<br>Mean ± SD 27.07 ± 4.64<br>Sample size (n) 6 mice<br>***Rheb<sup>CA</sup>+cGFP***<br>Mean ± SD 25.05 ± 3.97<br>Sample size (n) 11 mice<br>***Rheb<sup>CA</sup>+c4E-BP1<sup>CA</sup>***<br>Mean ± SD 27.69 ± 4.41<br>Sample size (n) 18 mice |
| **4E**<br>Bandpower, light<br>Theta | **One-way ANOVA**<br>p value 0.0408<br>F (DFn, DFd) F (2, 32) = 3.543 | ***GFP***<br>Mean ± SD 33.92 ± 2.84<br>Sample size (n) 6 mice<br>***Rheb<sup>CA</sup>+cGFP***<br>Mean ± SD 30.91 ± 2.62<br>Sample size (n) 11 mice<br>***Rheb<sup>CA</sup>+c4E-BP1<sup>CA</sup>***<br>Mean ± SD 32.89 ± 2.22<br>Sample size (n) 18 mice |
| **4E**<br>Bandpower, light<br>Alpha | **One-way ANOVA**<br>p value 0.7792<br>F (DFn, DFd) F (2, 32) = 0.251 | ***GFP***<br>Mean ± SD 14.25 ± 1.50<br>Sample size (n) 6 mice<br>***Rheb<sup>CA</sup>+cGFP***<br>Mean ± SD 14.35 ± 1.28<br>Sample size (n) 11 mice<br>***Rheb<sup>CA</sup>+c4E-BP1<sup>CA</sup>***<br>Mean ± SD 13.99 ± 1.44<br>Sample size (n) 18 mice |
| **4E**<br>Bandpower, light<br>Beta | **One-way ANOVA**<br>p value 0.0205<br>F (DFn, DFd) F (2, 32) = 4.398 | **GFP**<br>Mean ± SD 16.38 ± 2.64<br>Sample size (n) 6 mice<br>***Rheb<sup>CA</sup>+cGFP***<br>Mean ± SD 19.41 ± 2.78<br>Sample size (n) 11 mice<br>***Rheb<sup>CA</sup>+c4E-BP1<sup>CA</sup>***<br>Mean ± SD 16.99 ± 2.11<br>Sample size (n) 18 mice |
| **4E**<br>Bandpower, light<br>Gamma | **One-way ANOVA**<br>p value 0.0261<br>F (DFn, DFd) F (2, 32) = 4.094 | **GFP**<br>Mean ± SD 6.60 ± 1.91<br>Sample size (n) 6 mice<br>***Rheb<sup>CA</sup>+cGFP***<br>Mean ± SD 8.34 ± 1.98<br>Sample size (n) 11 mice<br>***Rheb<sup>CA</sup>+c4E-BP1<sup>CA</sup>***<br>Mean ± SD 6.57 ± 1.41<br>Sample size (n) 18 mice |
| **4F**<br>Bandpower, dark<br>Delta | **One-way ANOVA**<br>p value 0.3869<br>F (DFn, DFd) F (2, 32) = 0.978 | **GFP**<br>Mean ± SD 25.45 ± 6.32<br>Sample size (n) 6 mice<br>***Rheb<sup>CA</sup>+cGFP***<br>Mean ± SD 22.15 ± 5.10<br>Sample size (n) 11 mice<br>***Rheb<sup>CA</sup>+c4E-BP1<sup>CA</sup>***<br>Mean ± SD 24.58 ± 5.20<br>Sample size (n) 18 mice |
| **4F**<br>Bandpower, dark<br>Theta | **One-way ANOVA**<br>p value 0.0679<br>F (DFn, DFd) F (2, 32) = 2.928 | **GFP**<br>Mean ± SD 32.85 ± 3.26<br>Sample size (n) 6 mice<br>***Rheb<sup>CA</sup>+cGFP***<br>Mean ± SD 32.65.99 ± 2.61<br>Sample size (n) 11 mice<br>***Rheb<sup>CA</sup>+c4E-BP1<sup>CA</sup>***<br>Mean ± SD 34.99 ± 2.71<br>Sample size (n) 18 mice |
| **4F**<br>Bandpower, dark<br>Alpha | **One-way ANOVA**<br>p value 0.2188<br>F (DFn, DFd) F (2, 32) = 1.594 | **GFP**<br>Mean ± SD 14.62 ± 1.88<br>Sample size (n) 6 mice<br>***Rheb<sup>CA</sup>+cGFP***<br>Mean ± SD 16.21 ± 2.26<br>Sample size (n) 11 mice<br>***Rheb<sup>CA</sup>+c4E-BP1<sup>CA</sup>***<br>Mean ± SD 14.92 ± 2.12<br>Sample size (n) 18 mice |
| **4F**<br>Bandpower, dark<br>Beta | **One-way ANOVA**<br>p value 0.1481<br>F (DFn, DFd) F (2, 32) = 2.029 | **GFP**<br>Mean ± SD 16.80 ± 2.99<br>Sample size (n) 6 mice<br>***Rheb<sup>CA</sup>+cGFP***<br>Mean ± SD 18.57 ± 2.90<br>Sample size (n) 11 mice<br>***Rheb<sup>CA</sup>+c4E-BP1<sup>CA</sup>***<br>Mean ± SD 16.42 ± 2.75<br>Sample size (n) 18 mice |
| **4F** | **One-way ANOVA** | **GFP** |
| Bandpower, dark Gamma | p value 0.2268<br>F (DFn, DFd) F (2, 32) = 1.555 | Mean ± SD 8.65 ± 2.80<br>Sample size (n) 6 mice<br>***Rheb<sup>CA</sup>+cGFP***<br>Mean ± SD 8.47 ± 1.91<br>Sample size (n) 11 mice<br>***Rheb<sup>CA</sup>+c4E-BP1<sup>CA</sup>***<br>Mean ± SD 7.41 ± 1.54<br>Sample size (n) 18 mice |
| **5D**<br>Seizure frequency | **Mann-Whitney U test (two-tailed)**<br>p value 0.0260<br>Sum of ranks 213.5, 221.5<br>Mann-Whitney U 50.50 | ***Rheb<sup>CA</sup>+cGFP***<br>Mean ± SD 7.8 ± 4.7<br>Median 10.3<br>Sample size (n) 11 mice<br>***Rheb<sup>CA</sup>+c4E-BP1<sup>CA</sup>***<br>Mean ± SD 3.4 ± 5.0<br>Median 0.3<br>Sample size (n) 18 mice |
| **5E**<br>Seizures vs. cell placement | **Spearman's rank-order correlation**<br>p value <0.0001<br>Spearman r 0.7683 | Number of XY pairs 29 |
| **5F**<br>Seizures vs. cell size | **Spearman's rank-order correlation**<br>p value 0.0756<br>Spearman r -0.3351 | Number of XY pairs 29 |
| **5G**<br>Seizures vs. HCN4 intensity | **Spearman's rank-order correlation**<br>p value <0.0001<br>Spearman r 0.7445 | Number of XY pairs 29 |

For all patch clamp data, the numbers of recorded neurons are from 3 BFP+cGFP, 5 *Rheb<sup>CA</sup>+cGFP*, and 4 *Rheb<sup>CA</sup>+c4E-BP1<sup>CA</sup>* mice. Significant ANOVAs were further analyzed using Tukey's or Bonferroni's post-hoc test, as indicated on the figure legends; significant post-hoc results are noted with stars on the graph.

### Juvenile expression of c4E-BP1^CA^ restores the excitability of Rheb^CA^ neurons

To elucidate the contribution of 4E-BP to mTORC1-induced neuronal alterations, we examined whether compensating for dysregulated 4E-BP1 activity by juvenile 4E-BP1^CA^ expression would rescue Rheb^CA^ neuron electrophysiological function. We co-expressed Rheb^CA^ with a conditional 4E-BP1^CA^ (c4E-BP1^CA^) or GFP (cGFP) plasmid, as well as a tamoxifen-inducible Cre recombinase plasmid, in mice by IUE. In the conditional plasmids, 4E-BP1^CA^ or GFP is preceded by a loxP-flanked stop cassette which suppresses their expression. Upon tamoxifen administration, Cre recombinase mediates the excision of the stop cassette, allowing for 4E-BP1^CA^ or GFP to be expressed. As a control, mice were electroporated with BFP, cGFP, and Cre recombinase. A tdTomato reporter was included in all groups. Tamoxifen was administered in juvenile mice from P12 to P16 to induce 4E-BP1^CA^ or GFP expression. Wholecell patch clamp recordings were obtained from L2/3 pyramidal neurons expressing Rheb^CA^+c4E-BP1^CA^, Rheb^CA^+cGFP, or BFP+cGFP (hereafter referred to as control) in acute brain slices from P23-32 mice (**Figure 2A, B**).

**Figure 2:**
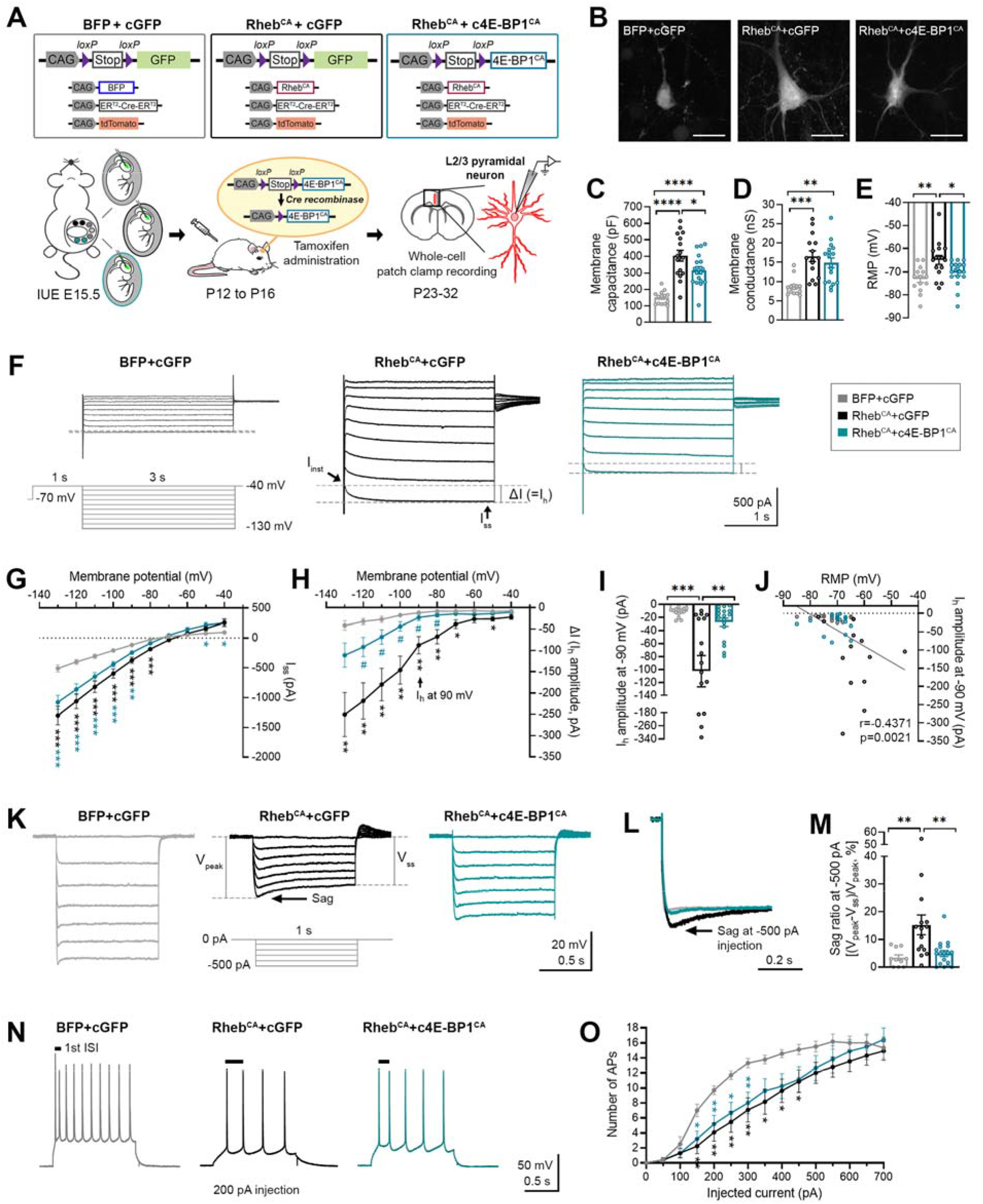
Juvenile expression of c4E-BP1^CA^ restores the excitability of Rheb^CA^ neurons. **(A)** Diagram of plasmids used to generate BFP+cGFP (control), Rheb^CA^+cGFP, a**nd** Rheb^CA^+c4E-BP1^CA^ mice and experimental strategy. Mouse embryos were electroporated **at** E15.5. For each litter, 1/3 of the embryos received BFP+cGFP, 1/3 received Rheb^CA^+cGFP, a**nd** 1/3 received Rheb^CA^+c4E-BP1^CA^ plasmids. A tamoxifen-inducible Cre (ER^T2^-Cre-ER^T2^) plasmid and a tdTomato reporter were co-electroporated in all 3 conditions. Tamoxifen was administered from P12 to P16 to induce GFP or 4E-BP1^CA^ expression. Whole-cell patch clamp recordings were performed in L2/3 pyramidal neurons in acute coronal slices from P23-32 mice. **(B)** Representative images of tdTomato+ (electroporated) neurons from brain slices used in patch clamp recording. The images are maximal intensity projections of 25 μm-thick z stacks sections. **(C-E)** Bar graphs of **(C)** membrane capacitance, **(D)** resting membrane conductance, and **(E)** resting membrane potential (RMP). **(F)** Representative current traces in response to a 1 s-long conditioning step to −40 mV from a holding potential of −70 mV, followed by a series of 3 s-long hyperpolarizing voltage steps from −130 mV to −40 mV in 10 mV increments. Traces at the −40 mV conditioning step are not shown due to overlapping traces from unclamped sodium spikes. *I_ss_, steady state current, I_inst_, instantaneous current*. **(G**) IV curve obtained from I_ss_ amplitudes. **(H)** IV curve obtained from I_h_ amplitudes or ΔI, where ΔI= I_ss_ -I_inst_. **(I)** Quantification of I_h_ amplitude at −90 mV. **(J)** Scatterplot of I_h_ amplitudes at −90 mV vs. RMP. n= 47 XY pairs. p=0.0021 by Pearson product-moment correlation. **(K)** Representative voltage traces in response to 1 s-long hyperpolarizing current steps from −500 pA to 0 pA in 100 pA increments from RMP. Arrow points I_h_-associated voltage sags induced by hyperpolarizing currents. **(L)** Representative voltage traces in response to a −500 pA current step. Traces were re-scaled and superimposed post-recording to visualize the differences in voltage sag size between groups. **(M)** Quantification of voltage sag ratio at −500 pA, where sag ratio = (V_peak_-V_ss_)/V_peak_ x 100. **(N)** Representative traces of action potential (AP) firing response to depolarizing current injections. **(O)** Input-output curve showing the mean number of APs in response to 500 ms-long depolarizing current steps from 0 to 700 pA in 50 pA increments. For all graphs, n=10-14 BFP+cGFP, 13-16 Rheb^CA^+cGFP, 17-18 Rheb^CA^+c4E-BP1^CA^ neurons. Data were analyzed using **(C-E, I, M)** one-way ANOVA with Tukey’s post-hoc test; *p<0.05, **p<0.01, ***p<0.001, ****p<0.0001 or **(G-H, O**) mixed-effects ANOVA with Tukey’s post-hoc test; *p<0.05, **p<0.01, ***p<0.001 (vs. BFP+cGFP), #p<0.05 (vs. Rheb^CA^+cGFP). Error bars are ± SEM.

Consistent with the visible changes in neuron size, Rheb^CA^+cGFP neurons displayed increased membrane capacitance which was partially rescued in Rheb^CA^+c4E-BP1^CA^ neurons (**Figure 2B, C, Table 1**). Rheb^CA^+cGFP neurons also had increased resting membrane conductance (**Figure 2D**) and more depolarized resting membrane potential (RMP) (**Figure 2E**). c4E-BP1^CA^ expression normalized the RMP but not the conductance (**Figure 2D, E, Table 1**).

We recently reported that Rheb^CA^ neurons have an abnormal expression of HCN4 channels, which give rise to a hyperpolarization-activated cation current (I_h_) that is normally absent in L2/3 pyramidal neurons (Hsieh et al., 2020). The aberrant I_h_, which has implications for neuronal excitability, preceded seizure onset and was detected by P8-12 in mice. Rapamycin treatment starting at P1 prevented aberrant HCN4 channel expression, suggesting their mTORC1-dependence. However, it is unknown whether HCN4 expression is regulated downstream through 4E-BPs and whether this electrophysiological alteration can be reversed. Thus, we examined the effects of c4E-BP1^CA^ expression on I_h_. To evoke I_h_, we applied a series of 3-second-long hyperpolarizing voltage steps from −130 mV to −40 mV. Consistent with previous findings (Hsieh et al., 2020), hyperpolarizing voltage pulses elicited significantly larger inward currents in Rheb^C^A+cGFP neurons compared to control neurons (**Figure 2F, G, Table 1)**. The inward currents in Rheb^CA^ neurons result from the activation of both inward-rectifier K+ channels (K_ir_) and HCN4 channels (Hsieh et al., 2020). Since K_ir_-mediated currents activate fast whereas I_h_ activates slowly during hyperpolarizing steps, to assess I_h_ amplitudes, we measured the difference between the instantaneous and steady-state currents at the beginning and end of the voltage pulses, respectively (i.e., ΔI) (Thoby-Brisson et al., 2000; Fan et al., 2016). The resulting IV curve revealed significantly larger I_h_ amplitudes in Rheb^CA^+cGFP neurons compared to control neurons where I_h_ was absent (**Figure 2H, Table 1**). These findings are best demonstrated at −90 mV, close to the reversal potential for K_ir_ channels, when no K_ir_ current is present and the observed current is predominantly due to I_h_ (**Figure 2I, Table 1**). c4E-BP1^CA^ expression did not affect the overall inward current amplitudes (**Figure 2F, G, Table 1)** but significantly decreased I_h_ amplitudes in Rheb^CA^ neurons (**Figure 2H, Table 1**). We did not detect changes in the overall inward current despite decreased I_h_ likely due to the small contribution of I_h_ to the overall inward current. At −90 mV, there were no differences in the mean I_h_ amplitudes between Rheb^CA^+c4E-BP1^CA^ neurons and control neurons, indicating c4E-BP1^CA^ expression restored aberrant I_h_ expression (**Figure 2I, Table 1**). Consistent with the function of I_h_ in maintaining RMP at depolarized levels (Lamas, 1998; Doan and Kunze, 1999; Lupica et al., 2001; Funahashi et al., 2003; Kase and Imoto, 2012), larger I_h_ amplitudes were significantly correlated with more depolarized RMPs in neurons (**Figure 2J, Table 1**). In agreement with the data for I_h_ amplitudes, hyperpolarizing current injections in current clamp mode evoked a prominent I_h_-mediated voltage sag in Rheb^CA^+cGFP neurons, which was absent or minimal in control and Rheb^CA^+c4E-BP1^CA^ neurons (**Figure 2K-M, Table 1**). Collectively, these findings demonstrate that c4E-BP1^CA^ expression rescues aberrant I_h_ expression in Rheb^CA^ neurons.

To assess the effects of c4E-BP1^CA^ expression on intrinsic excitability, we examined the action potential (AP) firing response to depolarizing current injections. We found no differences in AP threshold, peak amplitude, and half-width between the groups (**Supplemental Figure 2A-C, Table S1**). Rheb^CA^+cGFP neurons fired fewer APs for current injections above 100 pA compared to control neurons. No differences in the AP input-output curve were found between Rheb^CA^+cGFP and Rheb^CA^+c4E-BP1^CA^ neurons, indicating that c4E-BP1^CA^ expression did not rescue depolarization-induced firing frequency (**Figure 2N, O, Table 1**). In terms of firing pattern, neurons in all groups displayed a regular-spiking pattern with spike-frequency adaptation. However, while an initial doublet was observed in control neurons, consistent with the expected firing pattern for L2/3 mPFC pyramidal neurons (Kroon et al., 2019), this was absent in Rheb^CA^+cGFP neurons and reflected by a notably longer 1^st^ interspike interval (ISI) (**Figure 2N)**. Quantification of ISIs in an adaptive spike train of ≥10 spikes showed a significant increase in the 1^st^ ISI in Rheb^CA^+cGFP neurons which was normalized in Rheb^CA^+c4E-BP1^CA^ neurons. In comparison, no changes in the 9^th^ ISI were observed between the groups (**Supplemental Figure 2D, E, Table S1**). Thus, c4E-BP1^CA^ expression rescues irregular firing patterns but not depolarization-induced firing frequency in Rheb^CA^ neurons.

### c4E-BP1^CA^ expression in juvenile Rheb^CA^ mice reduces neuronal cytomegaly and aberrant HCN4 channel expression

Having established that c4E-BP1C^A^ expression corrects multiple electrophysiological deficits in Rheb^CA^ neurons, we next examined whether c4E-BP1^CA^ expression would also reverse mTORC1-induced histopathological abnormalities. We performed IUE with the same conditions as described above and administered tamoxifen from P28 to P32 to induce c4E-BP1^CA^ or cGFP expression in Rheb^CA^ mice (**Figure 3A**). By this age, FMCD pathology, including neuron misplacement and cytomegaly, is fully established and mice exhibit robust seizure activity (Lin et al., 2016; Zhang et al., 2020b). Brain tissue was collected at P84-106 (12-15 weeks) for histological examination.

**Figure 3:**
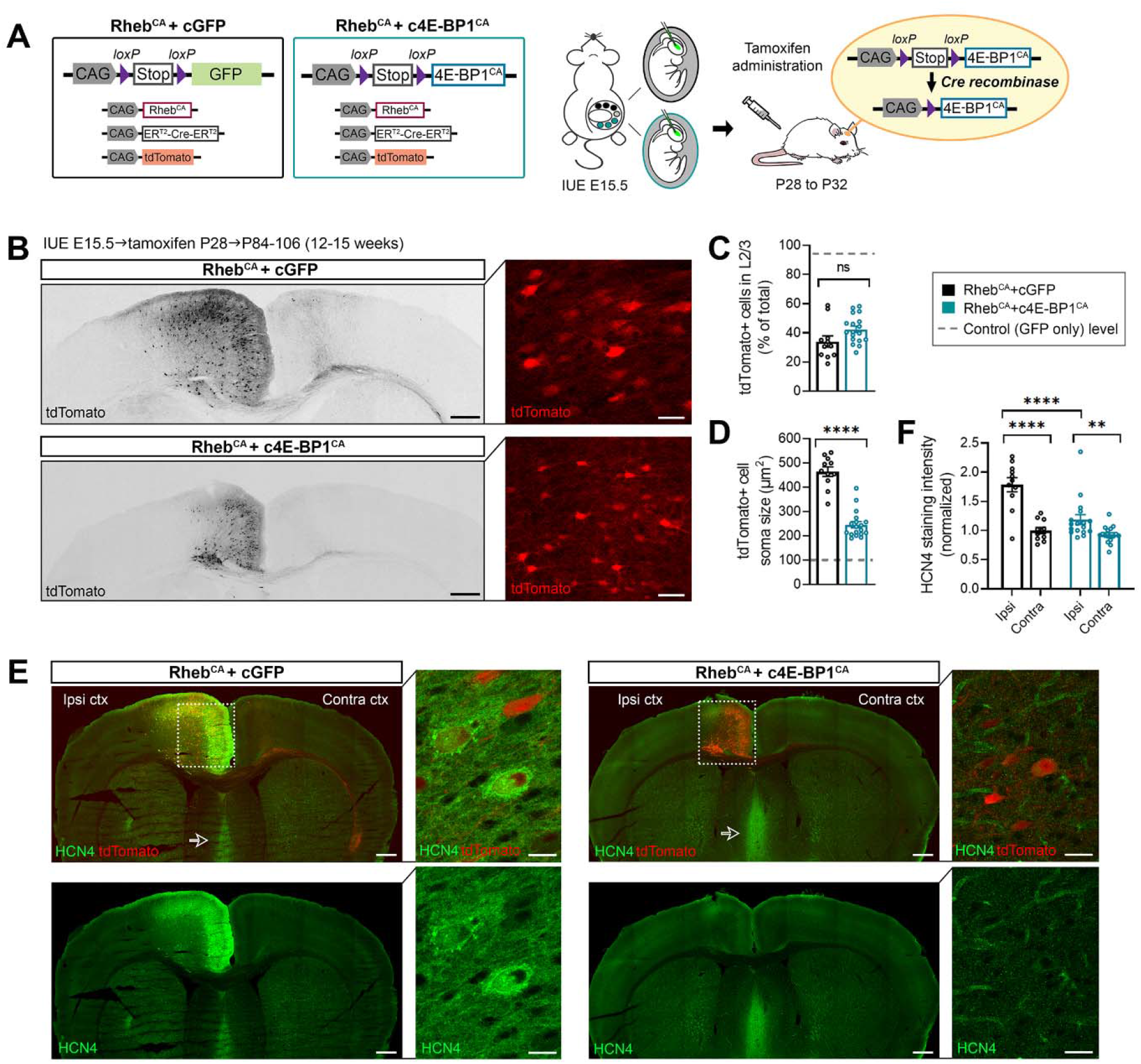
c4E-BP1^CA^ expression in juvenile Rheb^CA^ mice reduces neuronal cytomegaly and aberrant HCN4 channel expression. **(A)** Diagram of plasmids used to generate Rheb^CA^+cGFP and Rheb^CA^+c4E-BP1^CA^ mice and experimental strategy. Mouse embryos were electroporated at E15.5. For each litter, half of the embryos received Rheb^CA^+cGFP and the other half received Rheb^CA^+c4E-BP1^CA^. Tamoxifen-inducible Cre (ER^T2^-Cre-ER^T2^) and a tdTomato reporter were co-electroporated in both conditions. Tamoxifen was administered from P28 to P32 to induce GFP or 4E-BP1^CA^ expression. **(B)** Representative images of tdTomato+ cells in coronal sections from P84-106 Rheb^CA^+cGFP and Rheb^CA^+c4E-BP1^CA^ mice. Low magnification tile scan images showing tdTomato+ cell distribution across the cortical layers are on the left. High magnification images showing differences in tdTomato+ cell size are on the right. Scale bars=500 μm (left), 25 μm (right). **(C)** Quantification of tdTomato+ cell placement in L2/3. n=11 Rheb^CA^+cGFP, 18 Rheb^CA^+c4E-BP1^CA^ mice; each data point represents averaged values from 3 brain sections per animal. Data were analyzed by unpaired t-test. **(D)** Quantification of tdTomato+ cell soma size. n=11 Rheb^CA^+cGFP, 18 Rheb^CA^+c4E-BP1^CA^ mice; each data point represents averaged values from 50 cells per animal. Data were analyzed by unpaired t-test; ****p<0.0001. For graphs **(C)** and **(D)**, the gray dotted lines display previously reported levels for age-matched control mice electroporated with GPF only (Nguyen *et al., Journal of Neuroscience* 2019, DOI: https://doi.org/10.1523/JNEUROSCI.2260-18.2019) for comparison. **(E)** Representative images of tdTomato+ cells (red) and HCN4 staining (green, pseudocolored) in coronal sections from P84-106 Rheb^CA^+cGFP and Rheb^CA^+c4E-BP1^CA^ mice. Low magnification tile scan images showing HCN4 staining in whole brain sections are on the left. High magnification images showing somatic HCN4 staining in Rheb^CA^+cGFP neurons and lack of HCN4 staining in Rheb^CA^+c4E-BP1^CA^ neurons are on the right. White square denotes the area targeted by IUE. Arrows point to HCN4 staining in the medial septum. Scale bars=500 μm (left), 25 μm (right). **(F)** Quantification of HCN4 staining intensity. n=11 Rheb^CA^+cGFP, 17 Rheb^CA^+c4E-BP1^CA^ mice; each data point represents averaged values from 2 brain sections per animal. Data were normalized to the mean control and analyzed using two-way repeated measures ANOVA with Bonferroni’s post-hoc test; **p<0.01, ****p<0.0001. Error bars are ± SEM. *Ipsi ctx, ipsilateral cortex; contra ctx, contralateral cortex*.

We first examined the effects of c4E-BP1^CA^ expression on core FMCD cytoarchitectural abnormalities. We found no changes in Rheb^CA^ neuron placement following c4E-BP1^CA^ expression. In both groups, only ~40% of the electroporated neurons were correctly placed in L2/3 (**Figure 3B, C, Table 1**). In contrast, c4E-BP1^CA^ expression significantly decreased the soma size of Rheb^CA^ neurons (**Figure 3B, D, Table 1**). The mean soma size in a group of Rheb^CA^+c4E-BP1^CA^ mice that received vehicle treatment was not different from that in Rheb^CA^+cGFP mice, validating that the c4E-BP1^CA^ plasmid did not have leaky expression and the effects on neuron size require Cre-mediated c4E-BP1^CA^ expression (**Supplemental Figure 3A, B, Table S1**). Taken together, these findings show that c4E-BP1^CA^ expression in juvenile Rheb^CA^ mice reduces neuronal cytomegaly but not misplacement.

Given that c4E-BP1^CA^ expression normalized aberrant I_h_, in Rheb^CA^^CA^ neurons, we sought to further evaluate HCN channel expression by immunostaining. There are four HCN isoforms in the brain; HCN1 and HCN2 are highly expressed in the cortex and hippocampus, whereas HCN3 and HCN4 are predominantly found in subcortical regions (Notomi and Shigemoto, 2004; Oyrer et al., 2019). Within the cortex, HCN1 and 2 are expressed in the deeper layer pyramidal neurons, while no functional HCN channels are found in L2/3 pyramidal neurons (Kroon et al., 2019). We found no differences in HCN1-3 staining between hemispheres or animal groups (**Supplemental Figure 4A-F, Table S1**). In contrast, there was a significant increase in HCN4 staining intensity on the ipsilateral (electroporated) side of the cortex compared to the contralateral side in Rheb^CA^+cGFP mice (**Figure 3E, F, Table 1**). The overall staining pattern was diffuse with some neurons exhibiting bright somatic staining (**Figure 3E)**. The increase in HCN4 staining intensity was significantly reduced by c4E-BP1^CA^ expression (**Figure 3E, F, Table 1**), suggesting that the c4E-BP1^CA^-mediated rescue of I_h_ result from decreased HCN4 channel expression. In both groups, strong HCN4 staining was present in the medial septum where these channels are normally expressed (Notomi and Shigemoto, 2004), serving as an internal staining control (**Figure 3E**). The reduction in cortical HCN4 staining intensity was not detected in vehicle-treated Rheb^CA^+c4E-BP1^CA^ mice, supporting that the effects on HCN4 channel expression result from c4E-BP1^CA^ expression (**Supplemental Figure 4G, H, Table S1**). Collectively, these findings demonstrate that c4E-BP1^CA^ expression in juvenile Rheb^CA^ mice decreases aberrant HCN4 channel expression, consistent with the functional reduction in I_h_.

### c4E-BP1^CA^ expression after epilepsy onset in juvenile Rheb^CA^ mice normalizes cortical spectral activity and alleviates seizures

Considering the electrophysiological and morphological rescue following c4E-BP1^CA^ expression, we examined whether c4E-BP1^CA^ expression in juvenile Rheb^CA^ mice would improve cortical EEG activity and reduce seizures in adults. For these studies, the same animals from Figure 3 were used. The majority of these mice (6/11 Rheb^CA^+cGFP, 14/18 Rheb^CA^+c4E-BP1^CA^ mice) exhibited visible tonic-clonic seizures prior to or during tamoxifen administration from P28 to P35, thus, epilepsy was already manifested by the time c4E-BP1^CA^ expression was induced. Continuous vEEG recording was performed 6-8 weeks post-tamoxifen administration, starting at P76-94, for 7 days to monitor cortical activity and seizure behavior.

To examine cortical activity, we characterized the background EEG activity using spectral power analysis. This quantitative EEG method uses nonbiased signal processing to decompose raw EEG waves into discrete frequency bands and provides a neurophysiological measure of brain state and network function (Welch, 1967; Schnitzler and Gross, 2005; Kim and Im, 2018). Data were analyzed during the light and dark cycles to account for any circadian variables and were compared to a control group expressing only GFP. Rheb^CA^+cGFP mice displayed a significant upward shift in the relative power spectrum compared to control mice during light cycle activity. The shift was primarily observed in the higher frequency range of beta and gamma activity. No differences were observed between Rheb^CA^+c4E-BP1^CA^ and control mice, suggesting that c4E-BP1^CA^ expression normalized relative spectral power (**Figure 4A, B, Table 1**). Consistent with these findings, relative bandpower analysis revealed increased beta activity in Rheb^CA^+cGFP mice, which was normalized to control levels in Rheb^CA^+c4E-BP1^CA^ mice. Gamma activity was also higher in Rheb^CA^+cGFP mice than Rheb^CA^+c4E-BP1^CA^ mice, though neither group was statistically different from control mice (**Figure 4D, E, Table 1**). No differences in the dark cycle relative power spectrum or bandpower were observed between the groups (**Figure 4A, C, F, Table 1).** Overall, these data demonstrate the presence of altered cortical spectral activity in Rheb^CA^ mice which is restored by juvenile c4E-BP1^CA^ expression.

**Figure 4:**
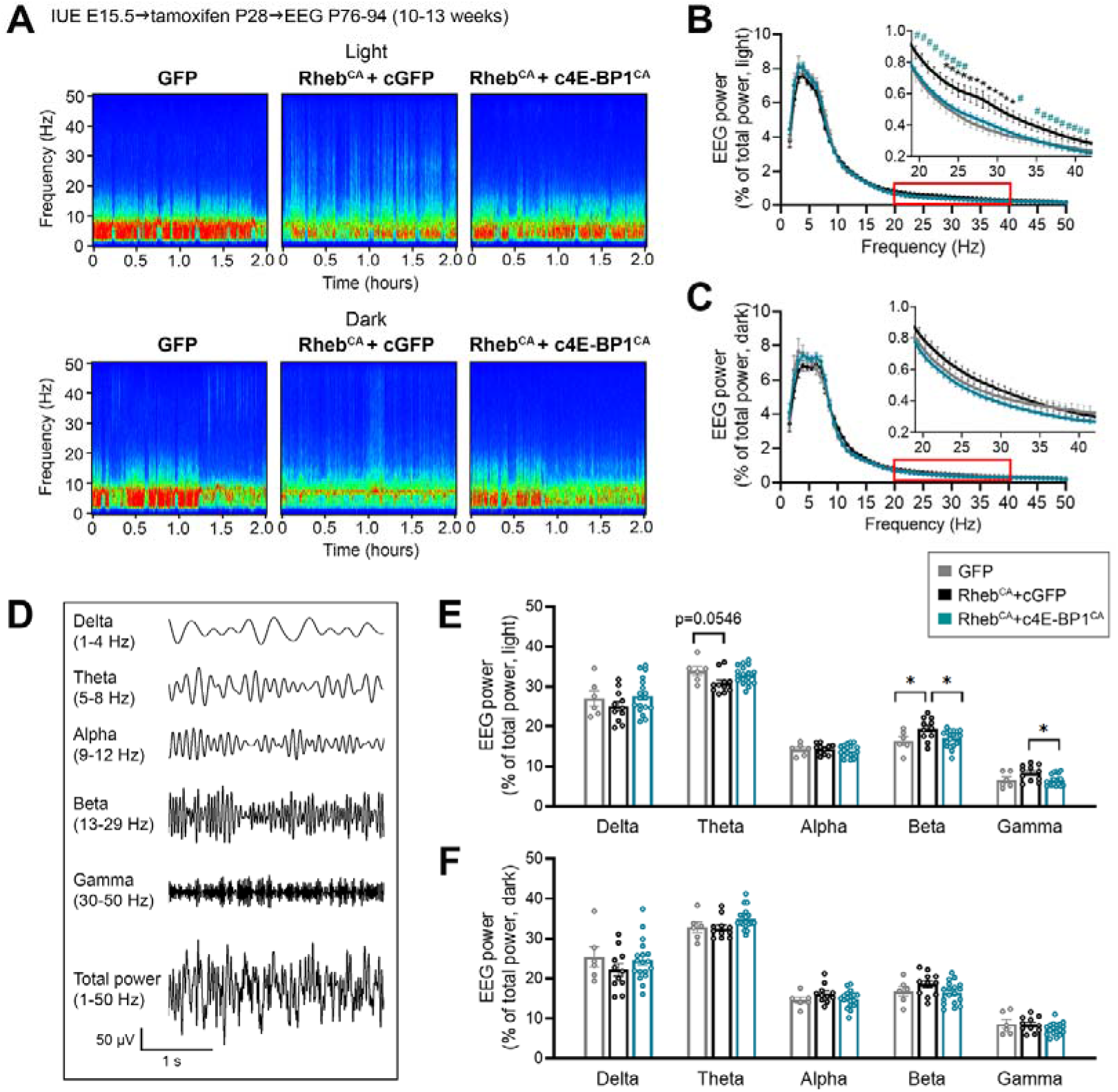
c4E-BP1^CA^ expression after epilepsy onset in juvenile Rheb^CA^ mice normalizes cortical spectral activity. **(A)** Representative spectrograms of background EEG during the light (12-2 pm, top) and dark (12-2 am, bottom) cycle. **(B, C)** Relative EEG power spectra of background EEG during the **(B)** light and **(C)** dark cycles. Insets show enlarged graphs from the area denoted with the red box. **(D)** Sample cortical EEG traces showing the delta, theta, alpha, beta, and gamma frequency bands. The bands were decomposed from the composite EEG signal shown at the bottom. **(E, F)** Bar graphs of the delta, theta, alpha, beta, and gamma relative bandpower during the **(E)** light and **(F)** dark cycles. For all graphs, n=6 GFP, 11 Rheb^CA^+cGFP, 18 Rheb^CA^+c4E-BP1^CA^ mice; each data point represents averaged values from 3-10 cycles (epochs) per animal. Data were analyzed using **(B, C)** two-way repeated measures ANOVA with Tukey’s post-hoc test; *p<0.05 (vs. GFP), #p<0.05 (vs. Rheb^CA^+cGFP) or **(E, F)** one-way ANOVA with Tukey’s post-hoc test; *p<0.05. Error bars are ± SEM.

We next quantified the incidence and frequency of seizures (**Figure 5A**). Comparing the portion of Rheb^CA^+c4E-BP1^CA^ mice to that of Rheb^CA^+cGFP mice, 44% vs. 9% had no electrographic seizures, 17% vs. 9% had <1 seizures/day, and 39% vs. 82% had >1 seizures/day (**Figure 5B, C**). The frequency of daily seizures was significantly decreased by >50% in Rheb^CA^+c4E-BP1^CA^ mice compared to Rheb^CA^+cGFP mice (**Figure 5D, Table 1**). Rheb^CA^+c4E-BP1 mice also had fewer seizures compared to vehicle-treated Rheb^CA^+c4E-BP1^CA^mice (**Supplemental Figure 5, Table S1**). Thus, these findings support that c4E-BP1^CA^ expression after epilepsy onset in juvenile Rheb^CA^ mice attenuates seizures.

**Figure 5:**
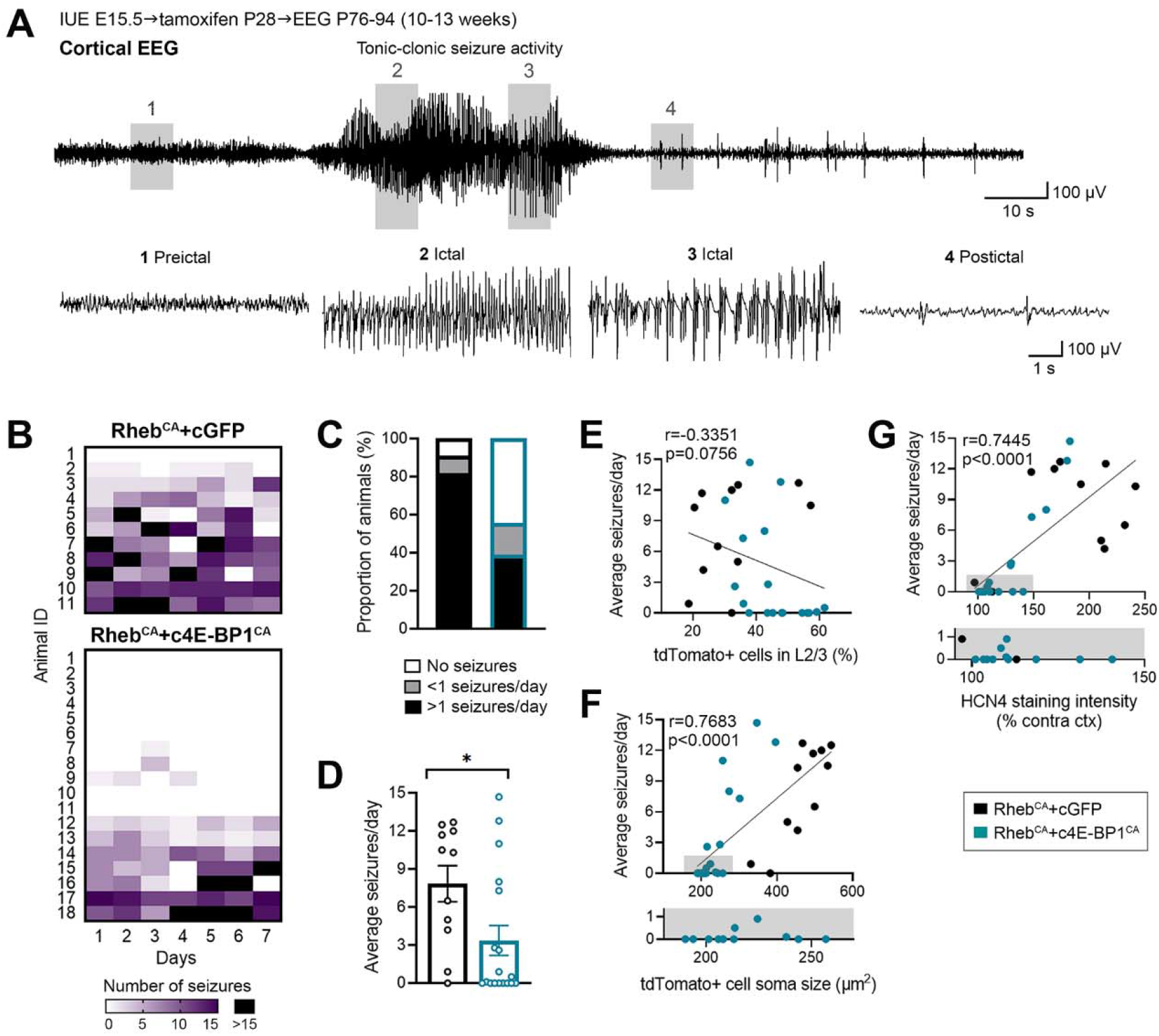
c4E-BP1^CA^ expression after epilepsy onset in juvenile Rheb^CA^ mice alleviates seizures. **(A)** Sample cortical EEG trace showing a typical seizure in Rheb^CA^ mice. Expanded EEG traces from representative preictal, ictal, and postictal periods are shown at the bottom. **(B)** Heatmap showing the daily number of seizures over 7 consecutive days for individual animals. **(C)** Bar graph showing the proportion of animals with no seizures, <1 seizure/day, and >1 seizure/day. **(D)** Quantification of seizure frequency. n=11 Rheb^CA^+cGFP, 18 Rheb^CA^+c4E-BP1^CA^ mice; each data point represents mean seizures/day from 7 days per animal. Data were analyzed using Mann-Whitney U test; *p=0.0206. **(E-G)** Scatterplots of **(E)** seizure frequency vs. cell placement in L2/3, **(F)** seizure frequency vs. cell size, and **(G)** seizure frequency vs. HCN4 staining intensity. For **(F)** and **(G)**, gray areas on the scatterplots are enlarged at the bottom of the respective plots to reveal overlapping values. n=29 **(E, F)**, 28 **(G)** XY pairs. Data were analyzed with Spearman’s rank-order correlation. Error bars are ± SEM.

Within both animal groups, we observed a broad range of seizure frequencies. To determine whether there was a relationship between individual animal seizure frequency and mTORC1-induced histopathological abnormalities, we correlated seizure frequency with the corresponding data for neuron placement, soma size, and HCN4 staining intensity as reported in Figure 3 for each animal. We found no correlation between seizure frequency and neuron placement (**Figure 5E, Table 1**). In contrast, seizure frequency positively correlated with soma size and HCN4 staining intensity (**Figure 5F, G, Table 1**). Based on these findings, we separated the HCN4 data from Figure 3 by seizure activity and found that Rheb^CA^+c4E-BP1^CA^ mice with <1 seizure/day had a complete rescue of HCN4 expression while those with >1 seizure/day had significantly elevated HCN4 levels (**Supplemental Figure 6, Table S1**). Taken together, these findings show that c4E-BP1^CA^-mediated reduction of seizure activity is associated with an improvement in neuron cytomegaly and aberrant HCN4 expression. Further, they show that c4E-BP1^CA^ expression attenuates seizures independently of neuron placement.

## DISCUSSION

Growing evidence supports mTORC1 pathway activating mutations as a major genetic cause for FMCD and intractable epilepsy, and understanding the mTORC1 downstream mechanisms that contribute to the pathology is crucial for developing new therapeutics. Here, we have identified the mTORC1-regulated translational repressor 4E-BP1 as a critical contributor to neuronal dysfunction and epilepsy. Our study is the first to demonstrate the reversibility of specific mTORC1-induced developmental alterations in neuron electrophysiological function and morphology by juvenile genetic manipulation of 4E-BP1 in mice. These findings have clinical relevance and support targeting 4E-BP1 as a potential treatment strategy for epilepsy in mTORC1-related FMCD.

Dysmorphic FMCD neurons from TSC and FCDII patients and Rheb^CA^ mice display increased levels of phosphorylated 4E-BP1, consistent with previous reports in human TSC, FCDII, and HME brain specimens (Baybis et al., 2004; Orlova and Crino, 2010; Hanai et al., 2017; Kim et al., 2019) and other FMCD mouse models with mTORC1 hyperactivation (Kassai et al., 2014; Kim et al., 2019; Koene et al., 2019). The presence of hyperphosphorylated 4E-BP1 implicate inhibited repressor activity, and thus, overactive translation. Recent work by Kim *et al*. reported increased translation of >200 mTOR-dependent genes with functions in epileptic seizures, dysmorphogenesis, and mislamination in cortical neurons from FMCD mice carrying brain somatic *mTOR* mutations (Kim et al., 2019). Reducing translation by eIF4E inhibition during fetal neurodevelopment rescued FMCD pathology and seizures, supporting dysregulated translation as a mechanism underlying FMCD and epilepsy (Kim et al., 2019). Our study is in agreement with these findings and further supports the role of translational dysregulation in epilepsy.

With regards to therapies for epilepsy, adult treatment with metformin, a widely used antidiabetic drug also known to inhibit mTORC1-mediated translation via AMP-activated protein kinase (AMPK) activation (Dowling et al., 2007), has been shown to decrease seizures in a mouse model of FMCD mice (Kim et al., 2019). However, since metformin activates multiple targets (Viollet et al., 2012) and has various effects in the brain, including inhibiting ERK signaling (Gantois et al., 2017), enhancing cerebral angiogenesis and neurogenesis (Wang et al., 2012; Zhu et al., 2020), decreasing glial activation and suppressing inflammatory pathways (Zhu et al., 2015; Ismaiel et al., 2016), and altering gut microbiome (Wu et al., 2017), it was unclear whether the mechanism of action was through inhibiting translation. In this study, we provide direct genetic support for reducing overactive translation via 4E-BP1 to alleviate established epilepsy. Given the recent advances in gene therapy for neurodevelopmental disorders (Deverman et al., 2018; Cheah et al., 2021; Turner et al., 2021), correcting translational dysregulation by 4E-BP1^CA^-targeted gene therapy may be a feasible treatment strategy for genetically defined FMCD.

Hyperactivation of mTORC1 signaling during neurodevelopment impairs cortical neuron migration and morphogenesis, resulting in neuronal misplacement across the cortical layer and cytomegaly. These FMCD hallmarks are consistently recapitulated in all current FMCD mouse models with mTORC1 hyperactivity (Nguyen and Bordey, 2021). Induction of c4E-BP1^CA^ expression at P28 in Rheb^CA^ mice was sufficient to reduce neuronal cytomegaly, but not misplacement. This latter finding is expected since pyramidal neurons reach their final position in the cortex by P7-10 (Greig et al., 2013; Kast and Levitt, 2019). At the behavioral level, c4E-BP1^CA^ expression improved cortical spectral activity and decreased seizures. Thus, our findings suggest that correcting or attenuating specific morphological and functional alterations alleviate FMCD-related epilepsy, even if the associated neuroanatomical defects are not reversed.

Few studies have examined the changes in intrinsic electrophysiological properties in cortical pyramidal neurons following mTORC1 hyperactivation, and even fewer studies have done this in the mPFC. We recently reported a set of electrophysiological alterations in mPFC Rheb^CA^ neurons which includes depolarized RMP, increased membrane conductance resulting from increased K_ir_ current and I_h_ due to abnormal HCN4 channel expression, and reduced firing frequency upon depolarizing current injections (Hsieh et al., 2020). Here, we recapitulated these results and additionally report changes in Rheb^CA^ neuron firing pattern characterized by loss of an initial doublet firing. c4E-BP1^CA^ expression restored RMP and the firing pattern and removed aberrant I_h_ in Rheb^CA^ neurons, suggesting that these alterations are 4E-BP1-dependent. In contrast, c4E-BP1^CA^ expression did not rescue the K_ir_ current or depolarization-induced firing frequency, suggesting that other processes contribute to these changes. The aberrant expression of HCN4 channels has notable implications for Rheb^CA^ neuron excitability and firing. Despite that these neurons have decreased depolarization-induced excitability, the aberrant presence of HCN4 channels drives firing through a cAMP-dependent process, and silencing HCN4 was sufficient to reduce seizure activity in mice (Hsieh et al., 2020). Thus, 4E-BP1^CA^-mediated removal of aberrant HCN4 channels potentially contributes to seizure reduction. Similar to Rheb^CA^ neurons, FMCD pyramidal neurons expressing other mTORC1 activating variants, including another Rheb variant, *Rheb Y35L*, as well as in *Tsc1* and *Depdc5* knockouts, also display decreased depolarization-induced firing responses (Ribierre et al., 2018; Goz et al., 2020; Onori et al., 2020). How these neurons lead to hyperexcitability and seizures is unknown. Our data suggest that aberrant acquisition of HCN4 channels may be a shared mode of abnormal excitability in these neurons. Nevertheless, whether aberrant HCN4 expression occurs in these other gene variants remains to be examined.

To our knowledge, no other studies have characterized the background EEG activity of FMCD mouse models by spectral analysis. EEG waves, reflecting rhythmic neuronal network activity in the brain, oscillate in various frequency bands that correlate with specific physiological functions and behavioral states (Buzsaki and Watson, 2012). Quantification of EEG activity by spectral power analysis has been used clinically to aid in the diagnosis and evaluation of cortical activity and disease severity, as well as assessing therapeutics, in neurocognitive and psychiatric disorders (Fonseca et al., 2006; Kanda et al., 2009; Chiarenza et al., 2016; Franko et al., 2016; Sidorov et al., 2017; Roche et al., 2019; Martinez et al., 2020). Remarkably, despite Rheb^CA^ being expressed in a relatively small fraction of neurons and a limited area of the cortex, this was sufficient to induce detectable changes in cortical spectral activity. In particular, we found increased beta activity in Rheb^CA^ mice which was restored by c4E-BP1^CA^ expression. Interestingly, altered resting-state functional connectivity with increased beta and gamma bands has been reported in human FCD (Jeong et al., 2014; Jin et al., 2015). Beta waves are associated with cortical activation and high arousal states, and excessive beta activity has been implicated in stress, anxiety, insomnia, and attention deficit hyperactivity disorder (Clarke et al., 2001; Perlis et al., 2001; Clarke et al., 2008; Engel and Fries, 2010). Therefore, increased beta activity in Rheb^CA^ mice could reflect the presence of these behaviors. Neuropsychiatric co-morbidities such as sleep disruption are common in human TSC and childhood epilepsy (Hunt, 1993; Hunt and Stores, 1994; Bruni et al., 1995; Tsai et al., 2019) and have been described in a TSC mouse model (Zhang et al., 2020a). Notably, alterations in Rheb^CA^ mice beta activity were only observed during the light cycle, when mice generally spend more time in rest and sleep, but not during the dark cycle, when mice are awake. This could hint at a disturbance in normal sleep activity in Rheb^CA^ mice, however, additional in-depth sleep analysis using EMG recording will be required to examine this.

Overall, our study demonstrates that although mTORC1 activates multiple molecular pathways, remarkably, restoring 4E-BP1 function is sufficient to improve many aspects of the disorder, including neuron cytomegaly, electrophysiological impairments, cortical spectral dysfunction, and epilepsy. Here, we did not examine the effects of c4E-BP1^CA^ expression on dendrite and synaptic function or axonal activity. Alterations in these parameters, which contribute to local or long-range alterations in connectivity and excitability, have been reported in FMCD neurons with mTORC1 hyperactivation (Chen et al., 2015; Gong et al., 2015; Lin et al., 2016; Ribierre et al., 2018; Onori et al., 2020; Zhong et al., 2021). Thus, it is possible that c4E-BP1^CA^ expression also impacts dendrite and axonal function, leading to attenuated epilepsy. Furthermore, while our data underscore that many alterations can be rescued by c4E-BP1^CA^ expression, it should not be inferred that dysregulated 4E-BP1 activity alone causes seizures. A recent study showed that ablation of 4E-BP2, but not 4E-BP1, reduces the threshold to chemoconvulsant-induced seizures in mice, although neither resulted in spontaneous seizures (Sharma et al., 2021). Interestingly, the effects on seizure threshold were specific to 4E-BP2 deletion in parvalbumin inhibitory neurons and not in other inhibitory or excitatory neurons in the hippocampus. Data from that study suggest that inhibiting 4E-BP1 or 4E-BP2 function in pyramidal neurons may not be sufficient to trigger seizures. This further supports the notion that a combination of molecular alterations downstream of mTORC1 is necessary to induce seizures, however, rescuing one molecular alteration is sufficient to alleviate the seizure burden.

In conclusion, we show that targeting 4E-BP1 after the underlying circuit abnormalities are established corrects multiple mTORC1-induced neuronal phenotypes and alleviates epilepsy. Our findings support the emerging notion that dysregulated translation is a major contributor to FMCD and epilepsy. Finally, targeting 4E-BP1 via gene therapy may represent a treatment strategy for epilepsy in mTORC1-related FMCD.

## METHODS

### Human tissue samples

Resected brain tissue samples (from the cortex) were obtained from individuals who underwent epilepsy surgery at Texas Children’s Hospital (Houston, TX, USA; 1 FCDII, 3 TSC) and Xiangya Hospital-Central South University (Changsha, China, 2 FCDII). One TSC sample was obtained from Tuberous Sclerosis Alliance. Patient pathological diagnosis, sex, and age are listed in **Table 2**. Tissue collection and use were approved by the Institutional Review Boards at Baylor College of Medicine and Yale University School of Medicine and the human ethical committee at Xiangya Hospital-Central South University. All samples were obtained directly from the operating room. Samples used for immunofluorescence staining were rapidly frozen in OCT and stored at −80°C until use. Frozen, unfixed tissue was cut into 15 μm-thick sections using a cryostat and mounted onto glass slides. Mounted sections were then fixed in ice-cold 4% paraformaldehyde (PFA) for 5 min, followed by 3 x 5 min rinses in phosphate buffered saline (PBS) immediately before immunofluorescence staining as described below. Samples used for immunohistochemistry were fixed in 10% formalin and stored at room temperature until use. Fixed samples were cut into 3-5 um-thick sections and stained using standard immunohistochemistry protocol.

**Table 2:** Information on TSC and FCDII patient tissue samples.

| Subject ID | Pathological diagnosis | Sex | Age |
| --- | --- | --- | --- |
| TSC (#1) | TSC | Unknown | Unknown |
| TSC (#2) | TSC | Female | 4 years |
| TSC (#3) | TSC | Female | 2 years |
| FCD (#1) | FCDII | Male | 7 years |
| FCD (#2) | FCDII | Female | 13 years |
| FCD (#3) | FCDII | Female | 5 years |

### Animals

All animal procedures were performed in accordance with Yale University Institutional Animal Care and Use Committee’s regulations. All experiments were performed on CD-1 mice (Charles River Laboratories) of either sex.

### Plasmid DNA

pCAG-Rheb^CA^ (also known as Rheb^S16H^) was a gift from Dr. Kentaro Hanada (Maehama et al., 2008). pCAG-TagBFP was a gift from Dr. Joshua Breunig (Breunig et al., 2012). pCAG-tdTomato (#83029) was previously generated in our lab (Pathania et al., 2012) and is available through Addgene. pCALNL-GFP (conditional GFP or cGFP, #13770), pCAG-ER^T2^Cre-ER^T2^ (#13777), and pCAG-GFP (#11150) were obtained from Addgene. pCALNL-4E-BP1^CA^ (conditional 4E-BP1^CA^ or c4E-BP1^CA^; also known as c4E-BP1^F113^A) was generated by excising GFP from the pCALNL-GFP backbone through sequential digestion with EcoRI and NotI and inserting 4E-BP1^CA^ between the EcoRI and NotI sites.

### *In utero* electroporation (IUE)

Timed-pregnant embryonic day (E) 15.5 mice were anesthetized with isoflurane, and a midline laparotomy was performed to expose the uterine horns. A DNA plasmid solution (1.5 μl) was injected into the right lateral ventricle of each embryo using a glass pipette. For **Figure 1** and **Supplemental Figure 1B, C**, a solution consisting of 2.5 μg/μl Rheb^CA^ (or BFP) + 1.0 μg/μl tdTomato was injected per embryo. For **Supplemental Figure 1D, E**, a solution of 3.5 μg/μl Rheb^CA^ (or tdTomato) + 1.0 μg/μl GFP was injected per embryo. For **Figures 2-5**, a solution of 3.0 μg/μl c4E-BP1^CA^ (or cGFP) + 2.5 μg/μl Rheb^CA^ + 0.4 μg/μl ER^T2^Cre-ER^T^ + 0.8 μg/μl tdTomato as injected per embryo. For each litter, half of the embryos were injected with the experimental condition and the other half received the respective control condition. **Figure 2** also included a control group that received 2.5 μg/μl BFP + 0.8 μg/μl tdTomato + 0.4 μg/μl ER^T2^Cre-ER^T2^ + 3.0 μg/μl cGFP per embryo. All plasmid solutions were diluted in water and contained 0.03% Fast Green dye to visualize the injection. A 5 mm tweezer electrode was positioned on the embryo head and 6x 42V, 50 ms pulses at 950 ms intervals were applied using a pulse generator (ECM830, BTX) to electroporate the plasmids into neural progenitor cells. Electrodes were positioned to target expression in the mPFC. The uterine horns were returned to the abdominal cavity and embryos were allowed to continue with their development. Pups were screened at postnatal day (P) 0 to verify successful electroporation to the targeted region via reporter gene expression on a fluorescence-enabled stereomicroscope (Olympus SZX16) before the downstream experiments.

### Tamoxifen administration

Tamoxifen (Cayman Chemical #13258) was dissolved in corn oil (Sigma) at a stock concentration of 10 mg/ml for 4-6 hours and stored at 4°C, protected from light. Tamoxifen was prepared fresh prior to each experiment. Mice were administered tamoxifen via intraperitoneal (i.p.) injections at a dose of 75 mg/kg once daily for 5 days. Mice were injected from P12 to P16 for patch clamp recording (**Figure 2)** and P28 to P32 for histology and EEG (**Figures 3-5**). Control groups receiving vehicle were included.

### Brain tissue fixation and immunofluorescence staining

Mice were deeply anesthetized with sodium pentobarbital (85□mg/kg i.p. injection) and perfused with ice-cold PBS followed by 4% PFA. Whole brains were dissected and post-fixed in 4% PFA for 2 hours and then cryoprotected in 30% sucrose for 24-48 hours at 4°C until they sank to the bottom of the tubes. Brains were serially cut into 50□um-thick coronal sections using a freezing microtome and stored in PBS + 0.01% sodium azide at 4°C until use.

For immunofluorescence staining, human (mounted) and mouse (free-floating) tissue sections were washed in PBS + 0.1% triton X-100 (PBS-T) for 2 x 10 min and permeabilized in PBS + 0.3% triton X-100 for 20-30 min. Sections were then incubated in blocking buffer (10% goat serum + 0.3% BSA + 0.3% triton X-100 in PBS) for 1-2 hours at room temperature. Mounted human tissue sections were incubated in primary antibodies (diluted in 5% goat serum + 0.3% BSA + 0.1% triton X-100 in PBS) for 1 day in a humidity chamber at room temperature. Free-floating mouse tissue sections were incubated in primary antibodies for 2 days at 4°C. Sections were then washed in PBS-T for 3 x 10 min and incubated in secondary antibodies for 2 hours at room temperature. Sections were then incubated in DAPI for 10 min, washed in PBS-T for 3 x 30 min, and rinsed in PBS before being mounted onto slides and coverslipped.

### Antibodies

The following primary antibodies were used: SMI-311 (BioLegend #837801, 1:200), p-4E-BP1 Thr37/46 (Cell Signaling Technology #2855, 1:200 for immunofluorescence, 1:300 for immunohistochemistry), p-4E-BP1/2/3-Thr45 (Bioss Antibodies bs-6421R, 1:200, 1:500), anti-HCN4 (Alomone Labs APC-052, 1:500), anti-HCN1 (APC-056, 1:500), anti-HCN2 (APC-030, 1:500), and anti-HCN3 (APC-057, 1:500). The following secondary antibody was used: goat anti-rabbit IgG Alexa Fluor Plus 488, 594, and 647 (Invitrogen A#32723, A#32740, and #A32733, respectively, 1:500).

### Microscopy and image analysis

Images were acquired using Zeiss LSM 880 [(**Figures 1-3**, and **Supplemental Figure 1B**] or Olympus Fluoview FV1000 [**Figures 3** (for analysis), **Supplemental Figures 1D, 3,** and **4**] confocal microscopes. Tile scan images were stitched together using Zeiss Zen Blue software. All image analyses were done using ImageJ software (NIH) and were performed blinded to experimental groups. Data were quantified using grayscale images of single optical sections unless noted otherwise. Representative images were prepared using Adobe Photoshop CC. All images meant for direct comparison were taken with the same settings and were uniformly processed.

For **Figure 1**, cell size was quantified by tracing the soma of tdTomato+ cells and measuring the area. p-4E-BP1 staining intensity was quantified by measuring the mean gray value within the same traced cells. For **Supplemental Figure 1**, p-4E-BP1/2/3 staining intensity was quantified by measuring the mean gray value within tdTomato+ or GFP+ cells. For each animal, 22-30 (**Figure 1)** or 12 (**Supplemental Figure 1**) randomly selected cells from 2 brain sections were measured. Each data point represents averaged values per animal. Staining intensities were normalized to the respective mean control. For **Figure 3** and **Supplemental Figure 3**, cell size was quantified by tracing the soma of tdTomato+ cells and measuring the area. Cell size was analyzed from maximum intensity projection images created from a 20 μm-thick z stack of optical sections taken at 2 μm increments. For each animal, 50 randomly selected tdTomato+ cells from 3 brain sections were measured. Each data point represents averaged values for each animal. Cell placement was quantified by counting all tdTomato+ cells within an 850 μm x 850 μm region of interest (ROI) surrounding the electroporated cortex. Cells within 300 μm from the pial surface were considered correctly located in L2/3 whereas cells outside that boundary were considered misplaced. For each animal, 3 brain sections were analyzed. Each data point represents averaged values for each animal. Data are shown as percent of total counted tdTomato+ cells. For **Figure 3** and **Supplemental Figure 4**, HCN1-4 staining intensity was quantified using images containing both the ipsilateral (electroporated) and contralateral cortices within the same plane of view. For each brain section, the mean gray value was measured in 3 randomly selected, non-overlapping 200 x 200 μm ROIs within the electroporated area and in 3 position-matched ROIs on the non-electroporated contralateral side. We analyzed 2 brain sections per animal for HCN4 and one brain section per animal for HCN1, 2, and 3; each data point represents averaged values from 6 (or 3) ROIs per animal. HCN1-4 staining intensities were normalized to the respective mean control (of Rheb^CA^+cGFP contralateral cortex).

### Acute slice preparation and whole-cell patch clamp recording

P23-32 mice were deeply anesthetized with isoflurane. Whole brains were rapidly dissected and immersed in ice-cold oxygenated (95% O_2_/5%CO_2_), high-sucrose cutting solution (in mM: 213 sucrose, 2.6 KCl, 1.25 NaH_2_PO_4_, 3 MgSO_4_, 26 NaHCO_3_, 10 Dextrose, 1 CaCl_2_, 0.4 ascorbate, 4 Na-Lactate, 2 Na-Pyruvate, pH 7.4 with NaOH). 300□um-thick acute brain slices containing the mPFC were cut using a vibratome (Leica VT1000). Slices were allowed to recover in a holding chamber with oxygenated artificial cerebrospinal fluid (aCSF, in mM: 124 NaCl, 3 KCl, 1.25 NaH_2_PO_4_, 1 MgSO_4_, 26 NaHCO_3_, 10 Dextrose, 2 CaCl_2_, 0.4 ascorbate, 4 Na-Lactate, 2 Na-Pyruvate, 300 mOsm/kg, pH 7.4 with NaOH) at 32°C for 45 min before returning to room temperature (25°C) where they were kept for 8-10 hours during the experiment.

Whole-cell current clamp and voltage clamp recordings were performed in a recording chamber at 28°C, using pulled borosilicate glass pipettes (4-7 MΩ resistance, Sutter Instrument) filled with an internal solution (in mM: 125 K-gluconate, 4 KCl, 10 HEPES, 1 EGTA, 0.2 CaCl_2_, 10 di-tris-phosphocreatine, 4 Mg-ATP, 0.3 Na-GTP, 280 mOsm/kg, pH 7.3 with KOH). Fluorescent (electroporated) neurons in the mPFC were visualized using epifluorescence on an Olympus BX51WI microscope with a 40X water immersion objective. Recordings were acquired using an Axopatch 200B amplifier and pClamp software (Molecular Devices). Data were filtered at 5 kHz and digitized with Digidata 1320 (Molecular Devices).

The resting membrane potential (RMP) was recorded within the first 10 seconds after achieving whole-cell configuration in current clamp mode while the cell was at rest without any holding current. The membrane capacitance and I_h_, were measured in voltage clamp mode. The membrane capacitance was calculated by dividing the membrane time constant by the average input resistance obtained from the current response to a 500 ms-long ±5 mV voltage step from −70 mV holding potential. I_h_ was activated by a 1 s-long conditioning step to −40 mV from a holding potential of −70 mV, followed by a series of 3 s-long hyperpolarizing voltage steps from −130 mV to −40 mV in 10 mV increments. I_h_ amplitudes (ΔI) were measured as the difference between the instantaneous current immediately following each test potential (I_inst_) and the steadystate current at the end of each test potential (I_ss_) (Thoby-Brisson et al., 2000; Fan et al., 2016). The resting membrane conductance, voltage sag, action potential (AP) properties were measured in current clamp mode. The resting membrane conductance was calculated using the average membrane potential change from 10x -500 pA hyperpolarizing current injections while the cell was at rest. The voltage sag was evoked by a series of 1 s-long hyperpolarizing currents from −500 pA to 0 pA in 100 pA increments from rest. The amplitude of the voltage sag in response to a −500 pA hyperpolarizing current step (sag ratio) was quantified using the equation (V_peak_-V_ss_)/V_peak_ x 100, where V_peak_ is the maximum voltage deflection and V_ss_ is the steady state voltage at the end of the hyperpolarizing pulse (George et al., 2009; Fan et al., 2016). The AP input-output curve was generated by injecting 500 ms-long depolarizing currents steps from 0 to 2000 pA in 50 pA increments. Only values from 0 to 700 pA were plotted due to the inactivation of Na+ channels in BFP+cGFP control neurons at >700 pA injections. The AP threshold, peak amplitude, and half-width were analyzed from averaged traces of 5-10 consecutive APs induced by a 50 ms-long minimal current (needed to induce the first AP) + 10 pA. The AP threshold was defined as the membrane potential at which the first derivative of an evoked action potential achieved 10% of its peak velocity (dV/dt). The AP peak amplitude was defined as the difference between the peak and baseline. The AP half-width was defined as the duration of the AP at the voltage halfway between the peak and baseline. The 1^st^ and 9^th^ interspike intervals (ISI) were calculated by measuring the time between the peaks of the 1^st^ and 2^nd^ spike and the 9^th^ and 10^th^ spike, respectively, using the first trace with ≥10 spikes. All data analysis was performed offline using pClamp software and exported to GraphPad Prism 8 software for graphing and statistical analysis.

### Video-Electroencephalography (vEEG) recording and analysis

#### Electrode implantation

Mice were implanted with prefabricated EEG headmounts (Pinnacle Technology, Inc. **#**8201-EEG) at 9-11 weeks of age. Mice were anesthetized with isoflurane and positioned on a stereotaxic frame using ear bars. A rostro-caudal midline incision was made in the skin to expose the skull surface. Four pilot holes (two bilateral holes 1 mm anterior to bregma and two bilateral holes 5 mm posterior to bregma, each 1.5 mm lateral to the sagittal suture) were tapped through the skull to the dura mater using a 23-gauge needle to accommodate the EEG headmount. The headmount was attached on top of the skull with superglue and four stainless steel screws (Pinnacle Technology, cat. no. 8209) were threaded into the pilot holes. Silver conductive paint was applied around the screw threads to ensure solid connections with the headmount. The entire implant was insulated using dental acrylic. Mice were allowed to recover in their home cage for at least 5 days before vEEG recording.

#### vEEG recording

Mice were vEEG recorded starting at 10-13 weeks of age, during which they were housed in individual recording chambers in a light-, temperature-, and humidity-controlled room with *ad libitum* access to food and water. Synchronous vEEG recording was acquired using a tethered three-channel EEG system (Pinnacle Technology, Inc. #8200-K1-iSE3) and Sirenia Acquisition software (Pinnacle Technology, Inc.). EEG data were acquired at a 400 Hz sampling rate and 1.0 Hz high-pass filtering. Mice were recorded 24 hours/day for at least 7 consecutive days. The average recording hours per animal was 179.8 ± 16.7 SD hours.

#### Seizure analysis

Seizure frequency was analyzed using Sirenia Seizure Basic software (Pinnacle Technology, Inc.). All analyses were performed blinded to experimental groups. The entire EEG traces were manually reviewed for the occurrence of seizures, defined as a sudden onset of high amplitude activity with a characteristic pattern of progressive frequency and amplitude increases over the course of the event lasting ≥10 s. Video data were visually inspected for behavioral correlates, including myoclonic jerks, tonic-clonic activity, convulsions, and loss of postural control (rearing and falling), and were used as secondary verification of seizures. The mean number of seizures per day (seizures/day) was obtained by dividing the total number of seizures by total recording hours and multiplied by 24 for each animal. The proportion of mice with 0, >1, or <1 seizures/day was determined by dividing the number of animals within the respective categories by the total number of animals for each group.

#### Spectral power analysis

Spectral power analysis of background EEG data was performed using LabChart v8 (AD Instruments). In addition to the traces obtained from the Rheb^CA^+cGFP and Rheb^CA^+c4E-BP1^CA^ mice, we analyzed traces from 6 control mice that were electroporated with GFP only, which were acquired as part of a previous study (Nguyen et al., 2019). EEG data from these animals were acquired under the same conditions, settings, and age. For each animal, a 2-hour epoch during each of the light (12-2 pm) and dark (12-2 am) cycles were analyzed. Any seizures or artifacts within these epochs were removed prior to analysis. EEG data were converted into frequency components using fast Fourier transformation (FTT) with an FTT size of 512 and a Hann (cosine-bell) window with 50% overlap. Relative power spectra were obtained by dividing power values within each frequency bin by total power. Relative bandpower in the delta δ (1–4 Hz), theta θ (5–8 Hz), alpha α (9–12 Hz), beta β (13–29 Hz), and gamma γ (30–50 Hz) bands were calculated by dividing the sum of power within each frequency range by total power. Data from time-of-day-matched epochs over multiple days were averaged for each animal (3-5 epochs for GFP mice, 5–10 epochs for Rheb^CA^+cGFP and Rheb^CA^+c4E-BP1^CA^ mice per light/dark cycle). Data are shown as relative power to reduce individual animal variability stemming from headcap implantation.

### Study design and statistical analysis

Human tissue samples from 3 individuals with TSC and 3 individuals with FCDII were evaluated for p-4E-BP1 staining intensity. Tissue from 5 control and 6 Rheb^CA^ mice were analyzed for p-4E-BP1 and p-4E-BP1/2/3 intensity at P30. 3 control and 3 Rheb^CA^ mice were analyzed for p-4E-BP1/2/3 intensity at P165. For patch clamp recording, 14 BFP+cGFP, 16 Rheb^CA^+cGFP, and 18 Rheb^CA^+c4E-BP1^CA^ neurons from 3 BFP+cGFP, 5 Rheb^CA^+cGFP and 4 Rheb^CA^+c4E-BP1^CA^ mice that received tamoxifen from P12 to 16, respectively, were recorded. Some results have fewer neuron numbers due to the loss of cell or membrane seal integrity during recording; the specific number of recorded cells for each result is indicated in **Tables 1** and **S1** and the figure legends. The average age of recording was P26.8 ± 3.2 SD (range: 23-32). For histology and vEEG studies, 11 Rheb^CA^+cGFP and 18 Rheb^CA^+c4E-BP1^CA^ mice that received tamoxifen from P28 to P32, as well as 3 vehicle-treated Rheb^CA^+c4E-BP1^CA^ (littermate with 6 of the tamoxifen-treated Rheb^CA^+c4E-BP1^CA^), were evaluated. The animals first underwent vEEG recording and brain tissue was collected afterward for histological examination. The average age of EEG recording start was 12.0 ± 1.0 SD weeks (range: 10.9-13.4 weeks) and the average age of tissue collection was 13.6 ± 1.3 SD weeks (range: 12.0-15.1 weeks). EEG data acquisition and analysis were performed completely blinded to experimental groups, as animal identity and experimental groups could only be determined afterward by evaluating fluorescence markers on prepared brain sections. Data for EEG, cell size, cell placement, and HCN4 (as well as HCN1-3) staining were from the same set of animals and are correlated in **Figure 5E-G**. One Rheb^CA^+c4E-BP1^CA^ mouse was excluded from HCN4 analysis because the hemispheres were split during tissue collection and it was not possible to analyze the ipsi- and contralateral sides within the same field of view. All data collection and analysis were performed blinded to experimental groups. Littermate controls were used for all experiments. Statistical analyses were performed using GraphPad Prism 9 software. The significance level was set at p< 0.05. The specific tests that were used, test results, and sample size (n, number of animals or neurons) are summarized in **Tables 1** and **S1** and described in the figure legends. Data are presented as mean ± standard error of the mean (SEM).

## ACKNOWLEDGEMENTS

This work was supported by NIH-NINDS R01 NS086329 (AB), NIH-NICHD F32 HD095567 (LHN), American Epilepsy Society Postdoctoral Fellowship (LHN), Yale Brown Coxe Postdoctoral Fellowship (LHN), and Swebilius Foundation Grant (LHN). We thank the Tuberous Sclerosis Alliance for providing the human TSC specimen.

## AUTHOR CONTRIBUTIONS

LHN performed *in utero* electroporation, histology, EEG recording, and data analysis, and prepared all statistical analyses and figures in the manuscript. YX performed the patch clamp recordings and related data analysis. TM contributed to some of the image analyses. LZ provided some of the human FCDII tissue samples and obtained the related immunohistochemistry data. TVL generated the conditional 4E-BP1^CA^ plasmid. HAB and AEA aided in EEG spectral analysis and related data interpretation. AEA provided the TSC and some of the FCDII human tissue samples. LHN and AB designed the studies and wrote the manuscript.

