## Supplemental information for "Genetic expression of 4E-BP1 in juvenile mice alleviates mTOR-induced neuronal dysfunction and epilepsy"

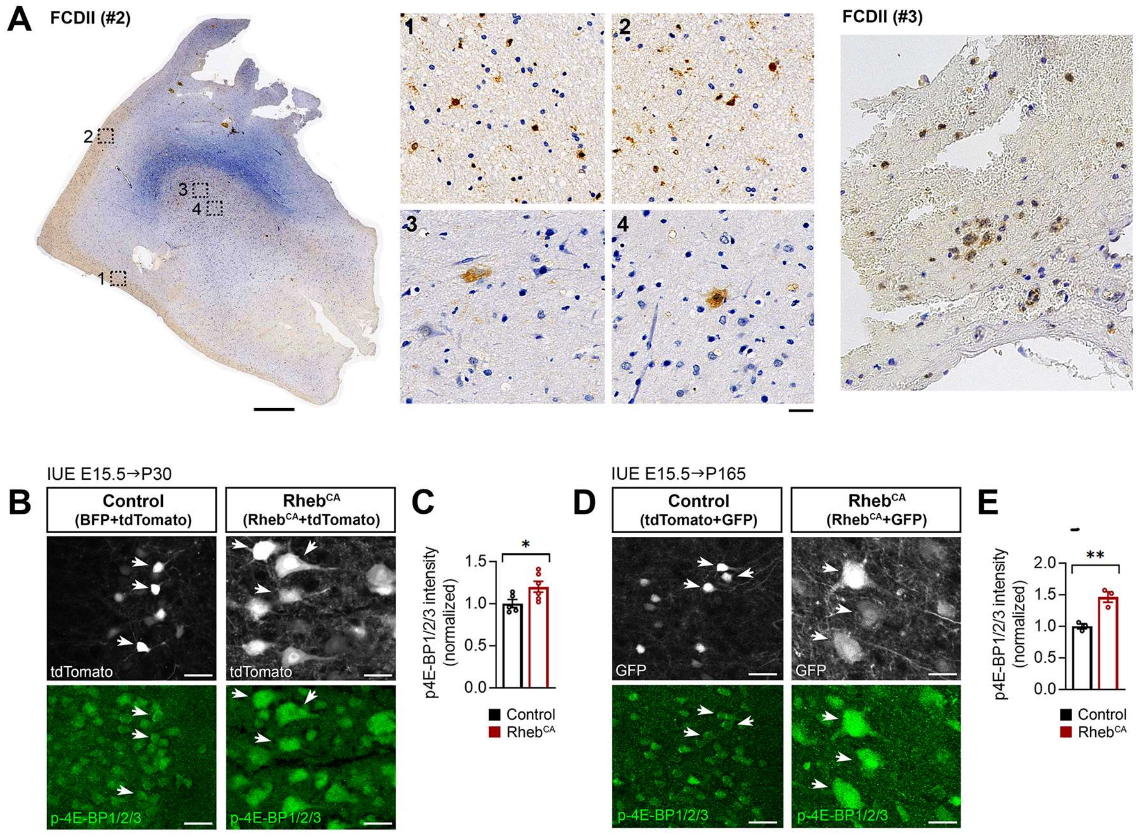


**Supplemental Figure 1 (related to Figure 1): Additional phospho-4E-BP1 and phospho-4E-BP1/2/3 immunostaining in FCDII human tissue and Rheb$^{CA}$ mice.**

**(A)** Representative images of phospho-4E-BP1 (p-4E-BP1) staining in resected brain tissue samples from 2 individuals with FCDII who underwent surgery for intractable epilepsy. Scale bars=1000 μm (left), 20 μm (right).

**(B)** Representative images of tdTomato+ cells (gray) and p-4E-BP1 staining (green, pseudocolored) in coronal cortical sections from P30 control and Rheb$^{CA}$ mice.

**(C)** Quantification of p-4E-BP1 intensity in tdTomato+ cells at P30. n=5 control, 6 Rheb$^{CA}$ mice; each data point represents averaged values from 12 cells per animal. Data were normalized to the mean control and analyzed by unpaired t-test; *p=0.0424.

**(D)** Representative images of GFP+ cells (gray) and p-4E-BP1 staining (green, pseudocolored) in coronal cortical sections from P165 control and Rheb$^{CA}$ mice.

**(E)** Quantification of p-4E-BP1 intensity in GFP+ cells at P165. n=3 control, 3 Rheb$^{CA}$ mice; each data point represents averaged values from 12 cells per animal. Data were normalized to the mean control and analyzed by unpaired t-test; *p=0.0071. Scale bars=25 μm.

Error bars are ± SEM.

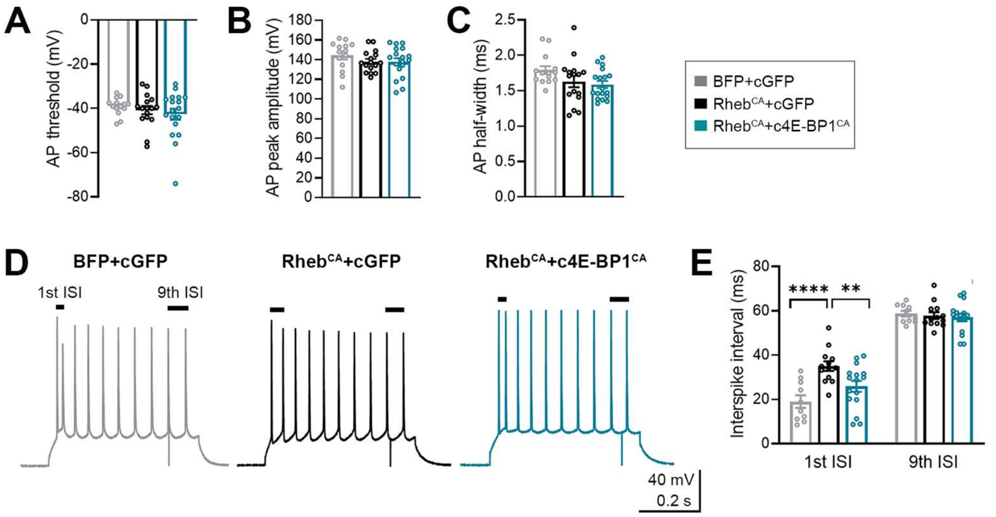


**Supplemental Figure 2 (related to Figure 2): Action potential properties and quantification of interspike intervals.**

**(A-C)** Bar graphs of action potential (AP) **(A)** threshold, **(B)** peak amplitude, and **(C)** half-width.

**(D)** Representative traces of AP firing response (with ≥10 spikes) to depolarizing current injections. The $1^{st}$ and $9^{th}$ interspike intervals (ISI) are denoted.

**(E)** Quantification of $1^{st}$ ISI and $9^{th}$ ISI.

**(A-C, E)** n=10-14 BFP+cGFP, 13-16 $Rheb^{CA}$+cGFP, 16-18 $Rheb^{CA}$+c4E-$BP1^{CA}$ neurons. Data were analyzed using **(A-C)** one-way ANOVA or **(E)** two-way repeated measures ANOVA with Tukey's post-hoc test. Error bars are ± SEM.

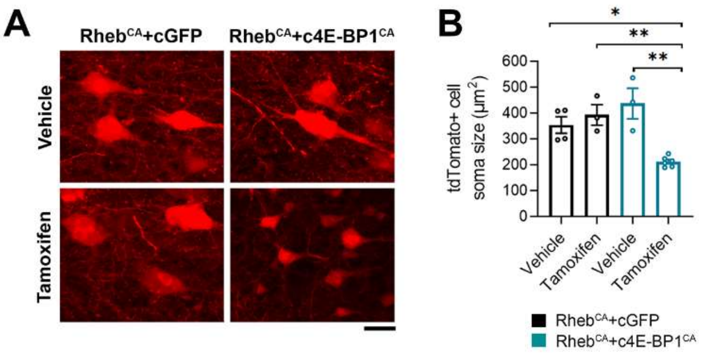


**Supplemental Figure 3 (related to Figure 3): Cortical neuron soma size in littermate vehicle- and tamoxifen-treated Rheb$^{CA}$+cGFP and Rheb$^{CA}$+c4E-BP1$^{CA}$ mice.**

Littermate Rheb$^{CA}$+cGFP or Rheb$^{CA}$+c4E-BP1$^{CA}$ mice were electroporated at E15.5 and treated with vehicle or tamoxifen from P28 to P32. Cell size analysis was performed at P84-106.

**(A)** Representative images of tdTomato+ cells in cortical sections from vehicle- and tamoxifen-treated Rheb$^{CA}$+cGFP and Rheb$^{CA}$+c4E-BP1$^{CA}$ mice.

**(B)** Quantification of tdTomato+ cell soma size. n=4 Rheb$^{CA}$+cGFP vehicle, 3 Rheb$^{CA}$+cGFP tamoxifen, 3 Rheb$^{CA}$+c4E-BP1$^{CA}$ vehicle, and 6 Rheb$^{CA}$+c4E-BP1$^{CA}$ tamoxifen mice; each data point represents averaged values from 50 cells per animal.

Data were analyzed by one-way ANOVA with Tukey's post-hoc test: *$p<0.05$, **$p<0.01$. Scale bars=25 μm. Error bars are ±SEM.

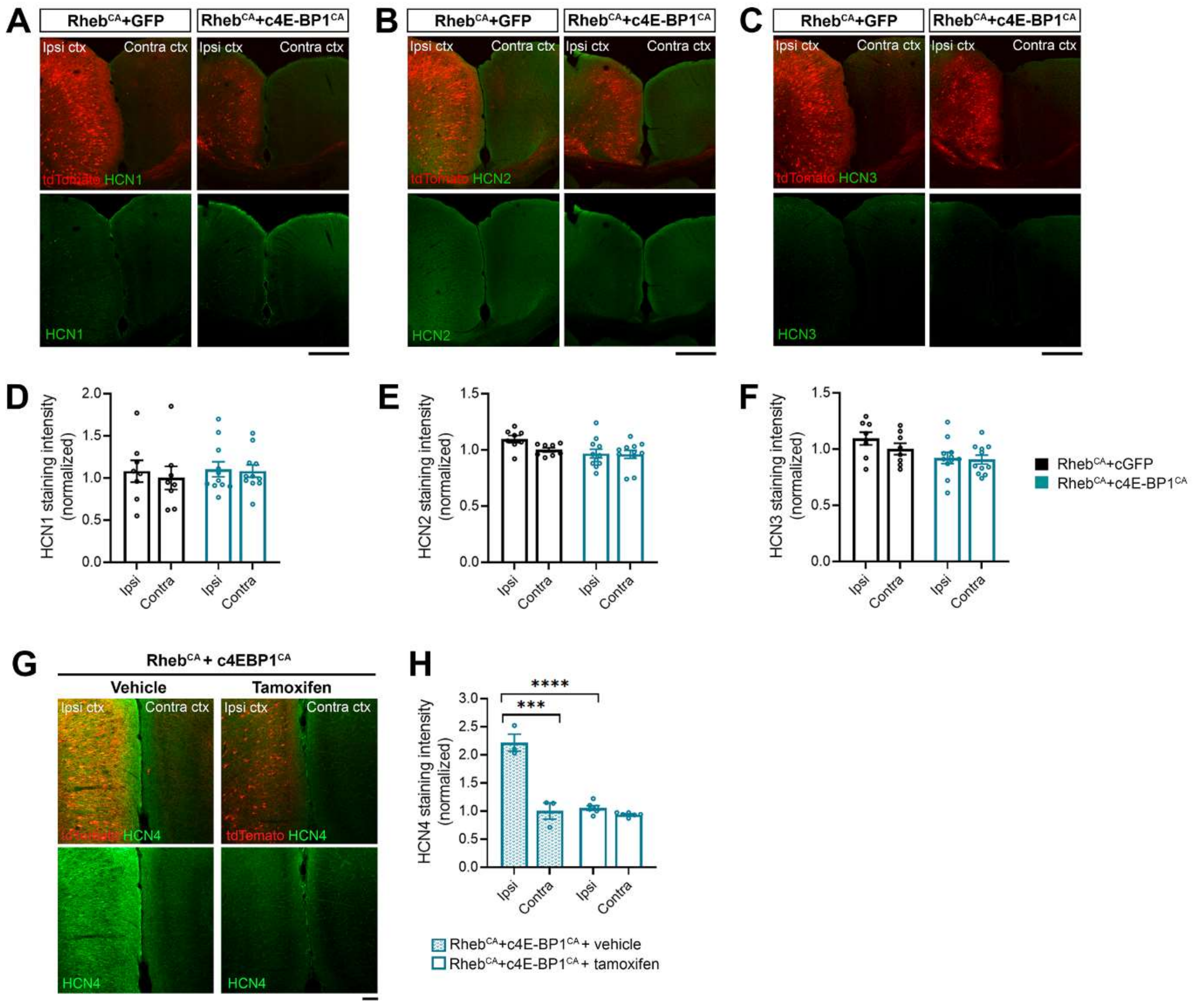


**Supplemental Figure 4 (related to Figure 3): HCN4 staining in littermate vehicle- and tamoxifen-treated Rheb$^{CA}$+c4E-BP1$^{CA}$ mice and HCN1-3 staining in Rheb$^{CA}$+cGFP and Rheb$^{CA}$+c4E-BP1$^{CA}$ mice.**

**(A-C)** Representative images of **(A)** HCN1, **(B)** HCN2, and **(C)** HCN3 staining in the cortex of Rheb$^{CA}$+cGFP and Rheb$^{CA}$+c4E-BP1$^{CA}$ mice. Merge images of tdTomato+ cells (red) and HCN1-3 staining (green, pseudocolored) are shown at the top. Single channel images of HCN1-3 staining are shown at the bottom. Scale bars=500 µm.

**(D-F)** Quantification of **(D)** HCN1, **(E)** HCN2, and **(F)** HCN3 staining intensity. n=8 Rheb$^{CA}$+cGFP, 11 Rheb$^{CA}$+c4E-BP1$^{CA}$ mice; each data point represents averaged values from 3 ROIs from one brain section per animal. Data were normalized to the mean and analyzed using two-way repeated measures ANOVA.

**(G)** Representative images of HCN4 staining in the cortex of littermate vehicle- and tamoxifen-treated Rheb$^{CA}$+c4E-BP1$^{CA}$ mice. Merge images of tdTomato+ cells (red) and HCN4 staining (green, pseudocolored) are shown at the top. Single channel images of HCN4 staining are shown at the bottom. Scale bar=100 µm.

**(H)** Quantification of HCN4 staining intensity. n=3 Rheb$^{CA}$+c4E-BP1$^{CA}$ vehicle, 6 Rheb$^{CA}$+c4E-BP1$^{CA}$ tamoxifen mice; each data point represents averaged values from 2 brain sections per animal. Data were normalized to the mean control and analyzed using two-way repeated measures ANOVA with Bonferroni's post-hoc test; ***p<0.001, ****p<0.0001.

Error bars are ± SEM. *Ipsi ctx, ipsilateral cortex; contra ctx, contralateral cortex.*

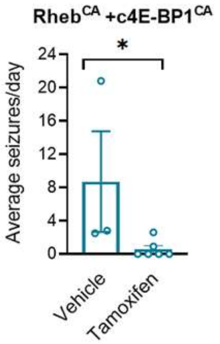


**Supplemental Figure 5 (related to Figure 5): Seizure frequency comparisons between vehicle- and tamoxifen-treated Rheb$^{CA}$+c4E-BP1$^{CA}$ mice.**

Quantification of seizure frequency in littermate vehicle- and tamoxifen-treated Rheb$^{CA}$+c4E-BP1$^{CA}$ mice. n=3 vehicle-treated, 6 tamoxifen-treated mice; each data point represents mean seizures/day from 7 days per animal. Data were analyzed using Mann-Whitney U test; *p=0.0238. Error bars are ± SEM.

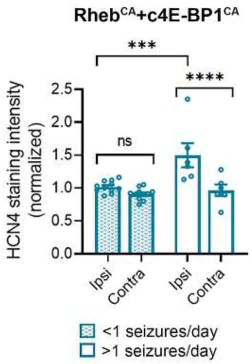


**Supplemental Figure 6 (related to Figures 3 and 5): HCN4 staining intensity comparisons between Rheb$^{CA}$+c4E-BP1$^{CA}$ mice with <1 and >1 seizures/day.**

Quantification of HCN4 staining intensity in Rheb$^{CA}$+c4E-BP1$^{CA}$ mice with <1 vs. >1 seizure/day. n=11 <1 seizure/day, 6 >1 seizure/day mice; each data point represents averaged values from 2 brain sections per animal. Data were normalized to the mean control (Rheb$^{CA}$+ cGFP contra ctx, as in Figure 3F) and analyzed using two-way repeated measures ANOVA with Bonferroni's post-hoc test; ***p<0.001, ****p<0.0001.

### Table S1: Table of supplemental results and statistics

| Figure | Statistical test, results | | Group mean, sample size | |
|---|---|---|---|---|
| **Supplemental 1C** | **Unpaired t-test (two-tailed)** | | ***Control (BFP)*** | |
| p-4E-BP1/2/3 intensity | p value | 0.0424 | Mean ± SD | 1.00 ± 0.11 |
| (P30) | t, df | t=2.363, df=9 | Sample size (n) | 5 mice |
| | | | ***Rheb$^{CA}$*** | |
| | | | Mean ± SD | 1.20 ± 0.16 |
| | | | Sample size (n) | 6 mice |
| **Supplemental 1E** | **Unpaired t-test (two-tailed)** | | ***Control (tdTomato)*** | |
| p-4E-BP1/2/3 intensity | p value | 0.0071 | Mean ± SD | 1.00 ± 0.07 |
| (P165) | t, df | t=5.082, df=4 | Sample size (n) | 3 mice |
| | | | ***Rheb$^{CA}$*** | |
| | | | Mean ± SD | 1.46 ± 0.14 |
| | | | Sample size (n) | 3 mice |
| **Supplemental 2A** | **One-way ANOVA** | | ***BFP+cGFP*** | |
| AP threshold | p value | 0.4666 | Mean ± SD | -39.0 ± 4.0 |
| | F (DFn, DFd) | F (2, 45) = 0.7753 | Sample size (n) | 14 neurons |
| | | | ***Rheb$^{CA}$+cGFP*** | |
| | | | Mean ± SD | -40.8 ± 7.6 |
| | | | Sample size (n) | 16 neurons |
| | | | ***Rheb$^{CA}$+c4E-BP1$^{CA}$*** | |
| | | | Mean ± SD | -42.7 ± 10.8 |
| | | | Sample size (n) | 18 neurons |
| **Supplemental 2B** | **One-way ANOVA** | | ***BFP+cGFP*** | |
| AP peak amplitude | p value | 0.3650 | Mean ± SD | 144.4 ± 15.0 |
| | F (DFn, DFd) | F (2, 45) = 1.031 | Sample size (n) | 14 neurons |
| | | | ***Rheb$^{CA}$+cGFP*** | |
| | | | Mean ± SD | 138.1 ± 11.0 |
| | | | Sample size (n) | 16 neurons |
| | | | ***Rheb$^{CA}$+c4E-BP1$^{CA}$*** | |
| | | | Mean ± SD | 137.9 ± 15.6 |
| | | | Sample size (n) | 18 neurons |
| **Supplemental 2C** | **One-way ANOVA** | | ***BFP+cGFP*** | |
| AP half-width | p value | 0.0792 | Mean ± SD | 1.79 ± 0.21 |
| | F (DFn, DFd) | F (2, 45) = 2.684 | Sample size (n) | 14 neurons |
| | | | ***Rheb$^{CA}$+cGFP*** | |
| | | | Mean ± SD | 1.63 ± 0.32 |
| | | | Sample size (n) | 16 neurons |
| | | | ***Rheb$^{CA}$+c4E-BP1$^{CA}$*** | |
| | | | Mean ± SD | 1.59 ± 0.21 |
| | | | Sample size (n) | 18 neurons |
| **Supplemental 2E** | **Two-way repeated measures ANOVA** | | ***BFP+cGFP*** | |
| 1st and 9th ISI | **ISI** | | Mean ± SD (1st ISI) | 18.95 ± 8.99 |
| | p value | <0.0001 | Mean ± SD (9th ISI) | 58.66 ± 4.08 |
| | F (DFn, DFd) | F (1, 36) = 314.1 | Sample size (n) | 10 neurons |
| | **Group** | | ***Rheb$^{CA}$+cGFP*** | |
| | p value | 0.0049 | Mean ± SD (1st ISI) | 35.02 ± 7.72 |
| | F (DFn, DFd) | F (2, 36) = 6.179 | Mean ± SD (9th ISI) | 57.82 ± 5.63 |
| | **ISI x group** | | Sample size (n) | 13 neurons |
| | p value | 0.0028 | ***Rheb$^{CA}$+c4E-BP1$^{CA}$*** | |
| | F (DFn, DFd) | F (2, 36) = 6.961 | Mean ± SD (1st ISI) | 25.80 ± 9.91 |
| | | | Mean ± SD (9th ISI) | 57.08 ± 6.93 |
| | | | Sample size (n) | 16 neurons |
| **Supplemental 3B** | **One-way ANOVA** | | ***Rheb$^{CA}$+cGFP vehicle*** | |

| | | | | |
|---|---|---|---|---|
| Cell soma size | **p value** | 0.0007 | Mean ± SD | 354.7 ± 63.7 |
| (P84-106) | **F (DFn, DFd)** | F (3, 12) = 11.78 | Sample size (n) | 4 mice |
| | | | ***Rheb$^{CA}$+cGFP tamoxifen*** | |
| | | | Mean ± SD | 393.7 ± 68.9 |
| | | | Sample size (n) | 3 mice |
| | | | ***Rheb$^{CA}$+c4E-BP1$^{CA}$ vehicle*** | |
| | | | Mean ± SD | 437.7 ± 102.5 |
| | | | Sample size (n) | 3 mice |
| | | | ***Rheb$^{CA}$+c4E-BP1$^{CA}$ tamoxifen*** | |
| | | | Mean ± SD | 212.6 ± 20.0 |
| | | | Sample size (n) | 6 mice |
| **Supplemental 4D** | **Two-way repeated measures ANOVA** | | ***Rheb$^{CA}$+cGFP*** | |
| HCN1 intensity | **Cortex side** | | Mean ± SD (ipsi) | 1.08 ± 0.37 |
| | p value | 0.0502 | Mean ± SD (contra) | 1.00 ± 0.39 |
| | F (DFn, DFd) | F (1, 17) = 4.442 | Sample size (n) | 8 mice |
| | **Group** | | ***Rheb$^{CA}$+c4E-BP1$^{CA}$*** | |
| | p value | 0.7222 | Mean ± SD (ipsi) | 1.10 ± 0.30 |
| | F (DFn, DFd) | F (1, 17) = 0.1306 | Mean ± SD (contra) | 1.08 ± 0.24 |
| | **Cortical side x group** | | Sample size (n) | 11 mice |
| | p value | 0.245 | | |
| | F (DFn, DFd) | F (1, 17) = 1.450 | | |
| **Supplemental 4E** | **Two-way repeated measures ANOVA** | | ***Rheb$^{CA}$+cGFP*** | |
| HCN2 intensity | **Cortex side** | | Mean ± SD (ipsi) | 1.10 ± 0.09 |
| | p value | 0.0668 | Mean ± SD (contra) | 1.00 ± 0.05 |
| | F (DFn, DFd) | F (1, 17) = 3.834 | Sample size (n) | 8 mice |
| | **Group** | | ***Rheb$^{CA}$+c4E-BP1$^{CA}$*** | |
| | p value | 0.0584 | Mean ± SD (ipsi) | 0.97 ± 0.13 |
| | F (DFn, DFd) | F (1, 17) = 4.119 | Mean ± SD (contra) | 0.96 ± 0.12 |
| | **Cortical side x group** | | Sample size (n) | 11 mice |
| | p value | 0.1045 | | |
| | F (DFn, DFd) | F (1, 17) = 2.942 | | |
| **Supplemental 4F** | **Two-way repeated measures ANOVA** | | ***Rheb$^{CA}$+cGFP*** | |
| HCN3 intensity | **Cortex side** | | Mean ± SD (ipsi) | 1.09 ± 0.16 |
| | p value | 0.0821 | Mean ± SD (contra) | 1.00 ± 0.14 |
| | F (DFn, DFd) | F (1, 17) = 3.415 | Sample size (n) | 8 mice |
| | **Group** | | ***Rheb$^{CA}$+c4E-BP1$^{CA}$*** | |
| | p value | 0.0526 | Mean ± SD (ipsi) | 0.92 ± 0.17 |
| | F (DFn, DFd) | F (1, 17) = 4.341 | Mean ± SD (contra) | 0.91 ± 0.13 |
| | **Cortical side x group** | | Sample size (n) | 11 mice |
| | p value | 0.1961 | | |
| | F (DFn, DFd) | F (1, 17) = 1.811 | | |
| **Supplemental 4H** | **Two-way repeated measures ANOVA** | | ***Rheb$^{CA}$+c4E-BP1$^{CA}$ vehicle*** | |
| HCN4 intensity | **Cortex side** | | Mean ± SD (ipsi) | 2.22 ± 0.27 |
| | p value | 0.0003 | Mean ± SD (contra) | 1.00 ± 0.25 |
| | F (DFn, DFd) | F (1, 7) = 43.49 | Sample size (n) | 3 mice |
| | **Group** | | ***Rheb$^{CA}$+c4E-BP1$^{CA}$ tamoxifen*** | |
| | p value | <0.0001 | Mean ± SD (ipsi) | 1.06 ± 0.10 |
| | F (DFn, DFd) | F (1, 7) = 258.3 | Mean ± SD (contra) | 0.94 ± 0.04 |
| | **Cortical side x group** | | Sample size (n) | 6 mice |
| | p value | 0.001 | | |
| | F (DFn, DFd) | F (1, 7) = 29.44 | | |
| **Supplemental 5** | **Mann-Whitney U test (two-tailed)** | | ***Rheb$^{CA}$+c4E-BP1$^{CA}$ + vehicle*** | |
| Seizure frequency | p value | 0.0238 | Mean ± SD | 8.7 ± 10.5 |
| | Sum of ranks | 23, 22 | Median | 2.8 |
| | Mann-Whitney U | 1 | Sample size (n) | 3 mice |

| | | |
|---|---|---|
| | | ***Rheb$^{CA}$+c4E-BP1$^{CA}$ + tamoxifen***<br>Mean ± SD 0.6 ± 1.1<br>Median 0.0<br>Sample size (n) 6 mice |
| **Supplemental 6**<br>HCN4 intensity | **Two-way repeated measures ANOVA**<br>**Cortex side**<br>p value <0.0001<br>F (DFn, DFd) F (1, 15) = 47.34<br>**Group**<br>p value 0.0161<br>F (DFn, DFd) F (1, 15) = 7.348<br>**Cortical side x group**<br>p value 0.0004<br>F (DFn, DFd) F (1, 15) = 20.35 | ***Rheb$^{CA}$+c4E-BP1$^{CA}$ <1 seizures/day***<br>Mean ± SD (ipsi) 1.02 ± 0.09<br>Mean ± SD (contra) 0.91 ± 0.09<br>Sample size (n) 11 mice<br>***Rheb$^{CA}$+c4E-BP1$^{CA}$ >1 seizures/day***<br>Mean ± SD (ipsi) 1.50 ± 0.45<br>Mean ± SD (contra) 0.97 ± 0.22<br>Sample size (n) 6 mice |

For all patch clamp data, the numbers of recorded neurons are from 3 BFP+cGFP, 5 Rheb$^{CA}$+cGFP, and 4 Rheb$^{CA}$+c4E-BP1$^{CA}$ mice. Significant ANOVAs were further analyzed using Tukey's or Bonferroni's post-hoc test, as indicated on the figure legends; significant post-hoc results are noted with stars on the graph.
